# A Futile Cycle? Tissue Homeostatic Trans-Membrane Water Co-Transport: Kinetics, Thermodynamics, Metabolic Consequences

**DOI:** 10.1101/2024.04.17.589812

**Authors:** Charles S. Springer, Martin M. Pike, Thomas M. Barbara

**Author notes:** **Corresponding author:** C. S. Springer;, Advanced Imaging Research Center, L452, Oregon Health and Science University, 3181 S. W. Sam Jackson Park Road, Portland, Oregon 97239.

## Abstract

The phenomenon of active trans-membrane water cycling (AWC) has emerged in little over a decade. Here, we consider H_2_O transport across cell membranes from the origins of its study. Historically, trans-membrane water transport processes were classified into: A) compensating bidirectional fluxes (“*exchange*”), and B) unidirectional flux (“*net flow*”) categories. Recent literature molecular structure determinations and molecular dynamic (MD) simulations indicate probably all the many different hydrophilic substrate membrane co-transporters have membrane-spanning hydrophilic pathways and co-transport water along with their substrates, and that they individually catalyze category A and/or B water flux processes, although usually not simultaneously. The AWC name signifies that, integrated over the all the cell’s co-transporters, the rate of *homeostatic*, bidirectional trans-cytolemmal water exchange (category A) is synchronized with the metabolic rate of the crucial Na^+^,K^+^-ATPase (NKA) enzyme. A literature survey indicates the stoichiometric (category B) water/substrate ratios of individual co-transporters are often very large. The MD simulations also suggest how different co-transporter reactions can be *kinetically* coupled molecularly.

Is this (Na^+^,K^+^-ATPase rate-synchronized) cycling futile, or is it consequential? Conservatively representative literature metabolomic and proteinomic results enable comprehensive free energy analyses of the many transport reactions with known water stoichiometries. Free energy calculations, using literature intracellular pressure (P_i_) values reveals there is an *outward* trans-membrane H_2_O barochemical gradient of magnitude comparable to that of the well-known *inward* Na^+^ electrochemical gradient. For most co-influxers, these gradients are finely balanced to maintain intracellular metabolite concentration values near their consuming enzyme Michaelis constants. The thermodynamic analyses include glucose, glutamate^**-**^, gamma-aminobutyric acid (GABA), and lactate^**-**^ transporters. 2%-4% P_i_ alterations can lead to disastrous concentration levels. For the neurotransmitters glutamate^**-**^ and GABA, very small astrocytic P_i_ changes can allow/disallow synaptic transmission. Unlike the Na^+^ and K^+^ electrochemical steady-states, the H_2_O barochemical *steady-state* is in (or near) chemical *equilibrium*. The analyses show why the presence of aquaporins (AQPs) does not dissipate the trans-membrane pressure gradient. A feedback loop inherent in the opposing Na^+^ electrochemical and H_2_O barochemical gradients regulates AQP-catalyzed water flux as an integral AWC aspect. These results also require a re-consideration of the underlying nature of P_i_. Active trans-membrane water cycling is not futile, but is inherent to the cell’s “NKA system” - a new, fundamental aspect of biology.

**SYNOPSIS:** *Via* intracellular pressure, membrane co-transported water influences thermodynamic control of cell metabolite maintenance.

## INTRODUCTION

One of the essential aspects of water in living tissue is its intra-extracellular compartmentalization. The kinetics of cellular, homeostatic trans-membrane water transport provides the basis of a new non-invasive, high-resolution MRI approach; metabolic activity diffusion imaging, MADI (1-3). It is providing novel views of cancer (4) and brain function (5) metabolism. Here, we inquire into kinetic, thermodynamic, and metabolic consequences of water co-transport phenomena in general.

### Distinct Trans-membrane Water Transport Processes

Category B *unidirectional* “flux” processes driven by trans-membrane osmotic gradients have been extensively investigated (6,7). These are characterized by cell volume (V) changes [cell swelling/shrinking; edema; tissue hypertrophy]. Their kinetics are quantified with the “osmotic” [flux, flow] water permeability coefficient, P_f_.

However, the earliest studies of which we are aware also detected category A *bidirectional* trans-membrane water “exchange” (8,9). In such processes, there is no V change – the systems are *homeostatic*. The kinetics are quantified with the “diffusional” water permeability coefficient, P_d_. Initial studies employed isotopic labeling (^2^HO^1^H), requiring very rapid, careful solution mixing – to perturb the system from isotopic equilibrium. As tracer methods, such approaches yield the P_d•_a_e_ product: a_e_ = ρ⟨A⟩, where ρ is the cell [number] density [cells/volume(tissue)] and ⟨A⟩ is the tissue-averaged cell surface area (1). [In the older literature, a_e_ is given as S, the *total* cell surface area per tissue [“ensemble”] volume.] P_d_ can be determined if a_e_ can be estimated. Early P_d_ values were compiled (10). We (11) and others showed that, at steady-state, P_d_ confounds V and A with the homeostatic cellular water efflux rate constant, k_io_, P_d_ = (V/A)k_io_. It is k_io_ that is a true measure of permeation probability (1). A nuclear magnetic resonance (NMR) approach was introduced in 1972 (12), which we later termed the “shutter-speed” (SS-NMR) method (13). Major SS-NMR advantages are it: **1.)** yields k_io_ un-confounded, and separated from (1 + ((ρ•V – 1)/f_W_)): f_W_ is the tissue water volume fraction (1,14); **2.)** does not require rapid solution mixing; and **3.)** could, in principle, quantify the category B flux and category A exchange processes simultaneously. An extracellular paramagnetic agent labels extracellular ^1^H_2_O magnetization, not extracellular water molecules (15), and non-invasive radio frequency electromagnetic pulses perturb the system magnetization from equilibrium. A compilation of early SS-NMR results was reported in (16). (Acronyms and symbols are listed in the **Appendix A1**).

### Molecular Aspects of Trans-membrane Water Transport

Over the years, many membrane-bound macromolecules have been found to transport water between intra- and extracellular spaces. In 1988, Agre and co-workers reported the selective water transport protein aquaporin (AQP) family (17). The impressive array of different AQP variants now known has been reviewed (6). During the 1990’s and 2000’s, many other proteins were found to co-transport water molecules, along with the metabolic substrates for which they are named (reviewed in (18,19)). **Figure 1** shows AQP and the ten water co-transporters detailed by Zeuthen (19). Each of their reactions is reversible, but here they are classified by their tendencies for influx or efflux under normal cellular conditions. These proteins all catalyze crucial processes, and are profitably considered as enzymes.

**Figure 1.**
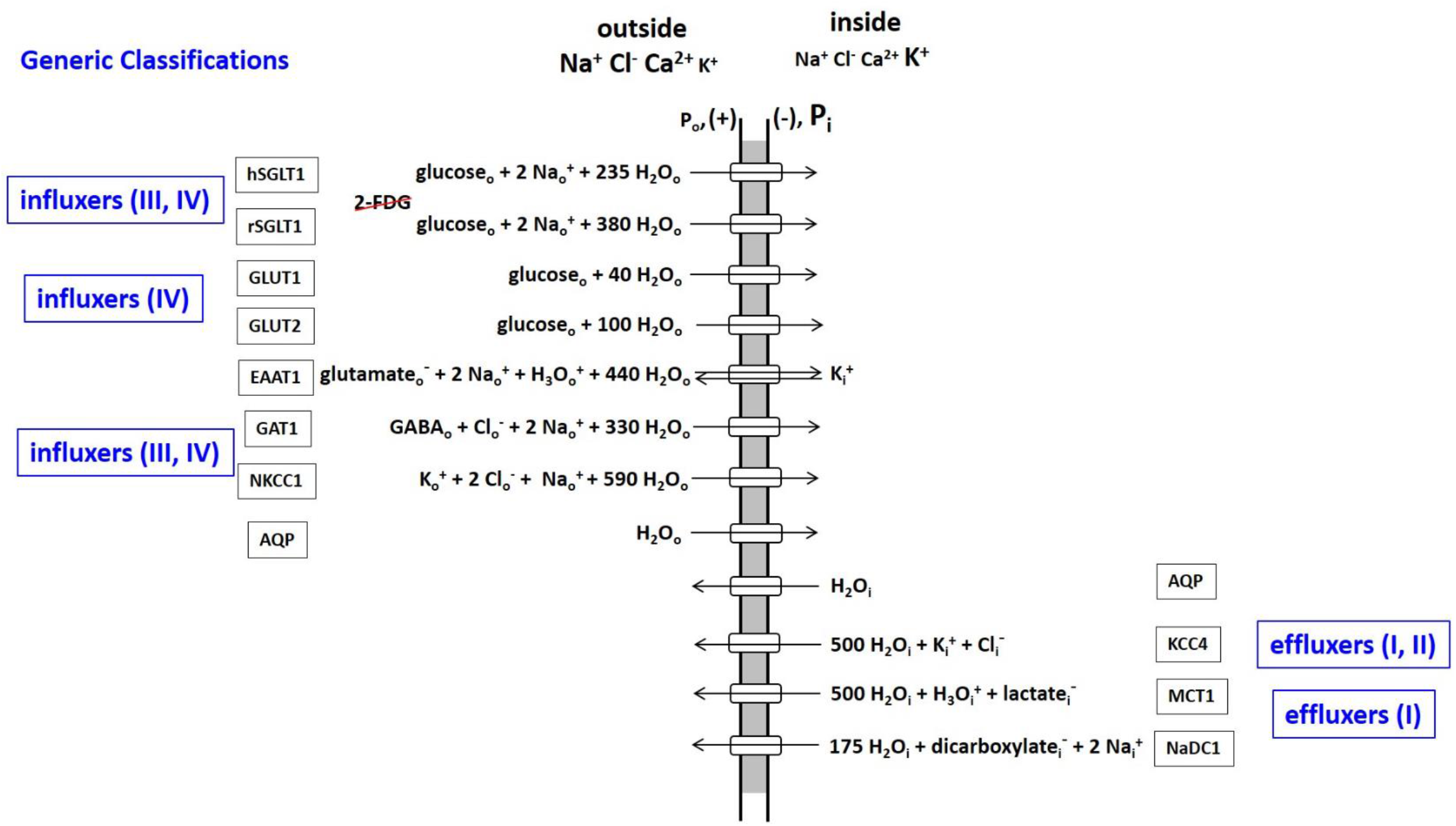
An Inventory of water transporting membrane proteins. The water stoichiometric values are taken from (19), and should be thought of as means of shot-to-shot variations (see text).

There are two glucose influx families: those co-transporting Na^+^ (SGLT), and those that do not (GLUT). (Interestingly, though both families transport glucose, it has been recently noted (20) the SGLT family does not transport 2-deoxy-2-^18^F-fluoro-D-glucose (2-FDG), the ^18^FDG-PET (positron emission tomography) glucose tracer. This can present interpretative problems.) The excitatory amino acid transporter (EAAT1) and the GABA transporter (GAT1) facilitate uptake of the principal excitatory and inhibitory neurotransmitters, glutamate^**-**^ and GABA [the gamma-aminobutyric acid zwitterion], respectively. (In the Fig. 1 EAAT1 influx reaction, note the H_3_O^+^ co-influx, and the K^+^ counter-*efflux* components (21).) The Na^+^,K^+^,2Cl^**-**^ co-transporter (NKCC1) facilitates K^+^ and Cl^**-**^ influx (22).

The K^+^,Cl^**-**^ co-transporter (KCC4) provides an important pathway for K^+^ and Cl^**-**^ efflux (22). The monocarboxylic acid transporter (MCT1) extrudes the lactate^**-**^ produced by cytoplasmic glycolytic-type metabolisms (HCO_3_^**-**^ is the analogous product of mitochondrial oxidative phosphorylation (23).

One is struck by the large water stoichiometries in most co-transporter cases. These values should be considered averages. Surely, they fluctuate stochastically (shot-to-shot) with each enzyme cycle (see below). Also, they likely vary with the cellular environment and biological condition. It is important to realize the Fig. 1 stoichiometries were determined in model systems *in vitro*, and by inducing net category B unidirectional fluxes (19). Homeostatic exchange was not studied in this regard.

In 1989, Ye and Verkman presented an elaborate optical method to measure P_f_ and P_d_ for *simultaneous* water efflux from and exchange in liposomes and in (adenosine tri-phosphate) ATP-free erythrocyte ghosts (24). Two very important results from that study are the following. First, a given water transporter can be involved in both category B flux and category A exchange processes, but not necessarily in the same proportions. Using HgCl_2_ to inhibit AQP, the authors found while AQP contributes 90% of ghost water efflux, it accounts for only 45% of ghost exchange flux. Second, though P_f_ for ghost efflux is numerically four times larger than P_d_ for ghost exchange (in the same units), the time-course for water efflux is 50 times longer. The exchange process is much, much faster. A confusing aspect is that P_f_ and P_d_ are always reported with the same dimensions (length/time). However, the two permeability coefficients are defined by rate laws of very different natures ((7); (10), pp. 44 ff), and are not at all the same. The P_f_ quantity characterizes transporter efficiency only in the presence of an osmotic gradient, and with the system perturbed from the steady-state. It gives no information on P_d_.

### Active Trans-Membrane Water Cycling (AWC)

During most *in vitro* and *in vivo* studies, the tissue is in homeostasis: there is no substantial cell swelling or shrinking. Only category A trans-membrane water bidirectional *exchange* obtains. Evidence mounts for a homeostatic, metabolic AWC process in cells (reviewed (1,2,15,25)). This is a very fast water molecule exchange, the kinetics of which are driven by, and synchronized to, those of the rate-limiting plasma membrane Na^+^,K^+^-ATPase (NKA) “sodium pump.” The AWC phenomenon was first detected using water proton (^1^H_2_O) SS-NMR in 2011 (26). **Figure 2** is a cartoon illustrating the AWC process. For each NKA cycle, one intracellular ATP_i_ molecule is hydrolyzed, three intracellular Na_i_^+^ ions expelled, and two extracellular K_o_^+^ ions imported. (First and second approximations of Fig 2 have appeared (2,15).) However, generic transporters II and III, respectively, allow K^+^ to re-exit and Na^+^ to re-enter exit the cell. Thus, NKA cycles extremely rapidly: depending on NKA expression, there can be 10^10^ (Na^+^ + K^+^) ions(cycled)/s/cell (15). These actions maintain the crucial trans-membrane ion concentration gradients, [Na_o_^+^] > [Na_i_^+^], [K_o_^+^] < [K_i_^+^]; and the membrane electrical potential (in more negative than out, Figs. 1,2). Explicit examples of II and III are KCC4 and SGLT, respectively. The NKA substrates intracellular ATP and Na^+^ and extracellular K^+^ (ATP_i_, Na_i_^+^, K_o_^+^, respectively) are rendered in red in Fig. 2, as is the natural product ouabain, a specific extracellular NKA inhibitor (ouabain_o_).

**Figure 2.**
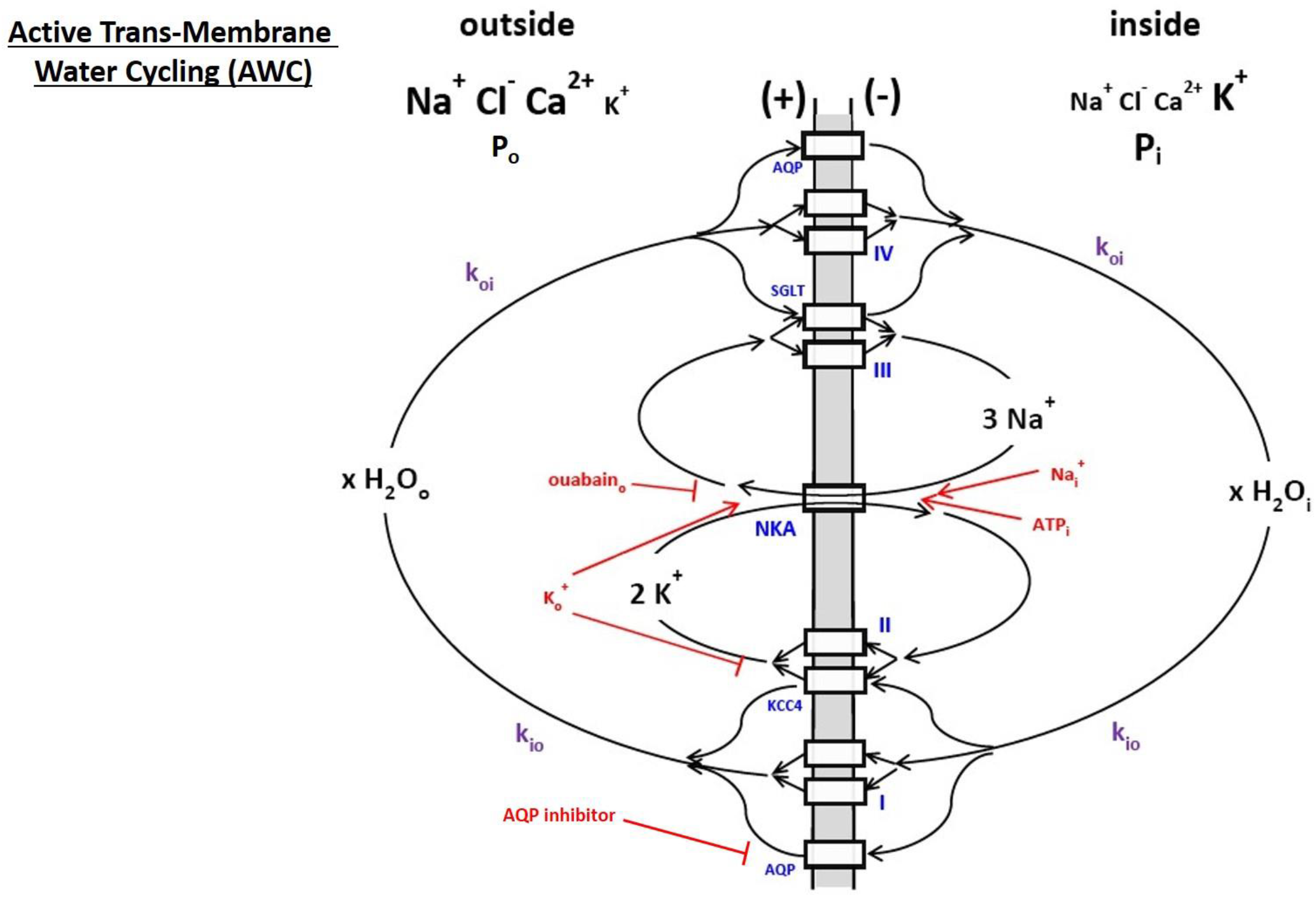
A cartoon of active trans-membrane water cycling (AWC). The rate constant for steady-state cellular water efflux is k_io_; that for steady-state cellular water influx is k_oi_. The Na^+^,K^+^-ATPase enzyme is indicated as NKA, while AQP, KCC4, and SGLT are defined in Figure 1. The actions of NKA substrates intracellular Na^+^ and ATP (Na_i_^+^ and ATP_i_) and extracellular K^+^ (K_o_^+^) are indicated in red, as are the NKA inhibitor extracellular ouabain (ouabain_o_) and an extracellular AQP inhibitor. Generic transporters I, II, III, and IV are exemplified in Figure 1: the actions of AQP, KCC4, and SGLT are shown here as specific examples. The quantity x is the AWC water stoichiometry integrated over the entire cell. Thus, for example, ^c^MR_H2O_(influx) = ^c^MR_H2O_(efflux) = x^c^MR_NKA_ = x(^c^MR_Na+_(influx))/3 = x(^c^MR_K+_(efflux))/2, where ^c^MR represents a cellular metabolic rate. First and second approximations of this cartoon have appeared in (15) and (2), respectively.

Generic water co-transporters I and IV represent *secondary, active* obligate water symporters for water to exit and enter cells, respectively. Most of the Fig. 1 enzymes fit into this classification: they share substrates (Na_o_^+^ and/or K_i_^+^) with NKA, a *primary, active* transporter (hydrolyzes ATP directly). The first-order, unidirectional rate constants k_io_ and k_oi_ are those for *cellular, homeostatic* water efflux and influx, respectively. In the differential first-order chemical trans-membrane water transport rate law, k_io_ can be expressed as the sum of energetically active, k_io_(a) and passive, k_io_(p), contributions that are further elaborated in **Equations (1)**, where: x is the overall, cellular AWC water

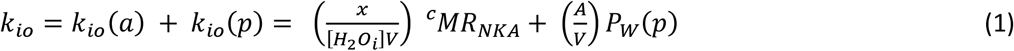

stoichiometric coefficient (H_2_O molecules(cycled)/NKA(cycle)/s/cell), [H_2_O_i_] the intracellular water concentration, ^c^MR_NKA_ the cellular NKA metabolic rate (ATP(molecules hydrolyzed by NKA)/s/cell), and P_W_(p) the non-metabolic, passive *diffusive* water membrane permeability coefficient P_d_(p) (1,2,15). The influx rate constant, k_oi_, is ρ-dependent, and also has active and passive components that need not be in the same proportions as for k_io_. In healthy, living tissue, k_io_(p) and k_oi_(p) seem negligible (2). There can be 10^12^ (H_2_O molecules actively cycled)/s/cell (15). The value of x may be of magnitude 10^6^ (2).

The sodium pump is perhaps biology’s most vital enzyme: it is found in all mammalian cells, and it’s role in the evolution of life is thought crucial (27). Since NKA homeostatic activity has never before been accessible *in vivo*, medical MR imaging (MRI) applications of MADI are very promising (1,2,4,5). The water proton MR signal (^1^H_2_O) is by far the largest from tissue (28). The AWC-based MADI approach has been deemed “a new paradigm” (3).

Water-selective AQP transporters can also be involved in AWC as long there is a thermodynamic driving force for water flux (our calculations suggest their role in cellular homeostasis, see Discussion). AQP molecules themselves comprise simple, *passive* channels (29,30).

The existence of AWC has been demonstrated by many different, deliberate manipulations of NKA kinetics with, for example, ATP_i_, K_o_^+^ and oubain_o_ concentration alterations (Fig. 2, red), while monitoring k_io_. More than 20 independent studies, of a range of tissues, have been itemized (2,25). AWC can be mapped *in vivo* with non-invasive MADI, which estimates k_io_, V, and ρ separately (1,2). In awake, resting human brain gray matter [GM], MADI mapping of the cellular water *flux* [H_2_O_i_]k_io_V = ^c^MR_AWC_, an x^c^MR_NKA_ estimate, correlates very well with the tissue metabolic rate of glucose consumption, ^t^MR_glu_ (glucose(consumed)/s/unit volume(tissue)) determined from quantitative ^18^FDG-PET (2). This is expected: overall ATP production (MR_ATP_) and consumption remain balanced in the resting brain. The white matter [WM] ^c^MR_AWC_/^t^MR_glu_ ratio is larger than the GM ^c^MR_AWC_/^t^MR_glu_ slope (2), likely indicating a more oxidative metabolic mechanism.

The most efficient MR_ATP_ comes from mitochondrial oxidative phosphorylation (Fig. 2 of ref (15)), and MR_AWC_ should be even more sensitive to this. Indeed, *in vitro* model brain tissue studies indicate k_io_V correlates strongly with mitochondrial MR_O2_(consumption) (31). This is shown in **Figure 3** in the context of MR_NKA_ stimulation *via* [K_o_^+^] titration (Fig. 2). The vertical axis measures the population-averaged ⟨k_io_⟩_n_ for organotypic, cultured [spiking] rat somatosensory cortex superfused with a paramagnetic agent. This SS-NMR study also allowed the estimation that ⟨V⟩_n_ was rather constant during the titration (Fig. 3D of ref. (31)): thus ⟨k_io_⟩_n_ was proportional to ⟨k_io_V⟩_n_ = ⟨^c^MR_AWC_⟩_n_. The horizontal axis measures ⟨^t^MR_O2_⟩_n_, directly determined in completely independent [K_o_^+^] titrations of isolated rat brain synaptosome suspensions. It is the [K_o_^+^] titrations that allow this correlation: these are shown as outer ordinate and abscissa scales. They are non-linearly (Michaelis-Menten) related to the inner ⟨k_io_⟩_n_ and ⟨^t^MR_O2_⟩_n_ scales, which are linear. The correlation is excellent over most of the range. (It is interesting O_2_ consumption continues in the synaptosome suspensions even when [K_o_^+^] is zero. This is not the case for k_io_ in the SS-NMR study of cultured cortex preparations.) The quantity k_io_V is probably best thought of as a high-resolution *in vivo* measure of mitochondrial function.

**Figure 3.**
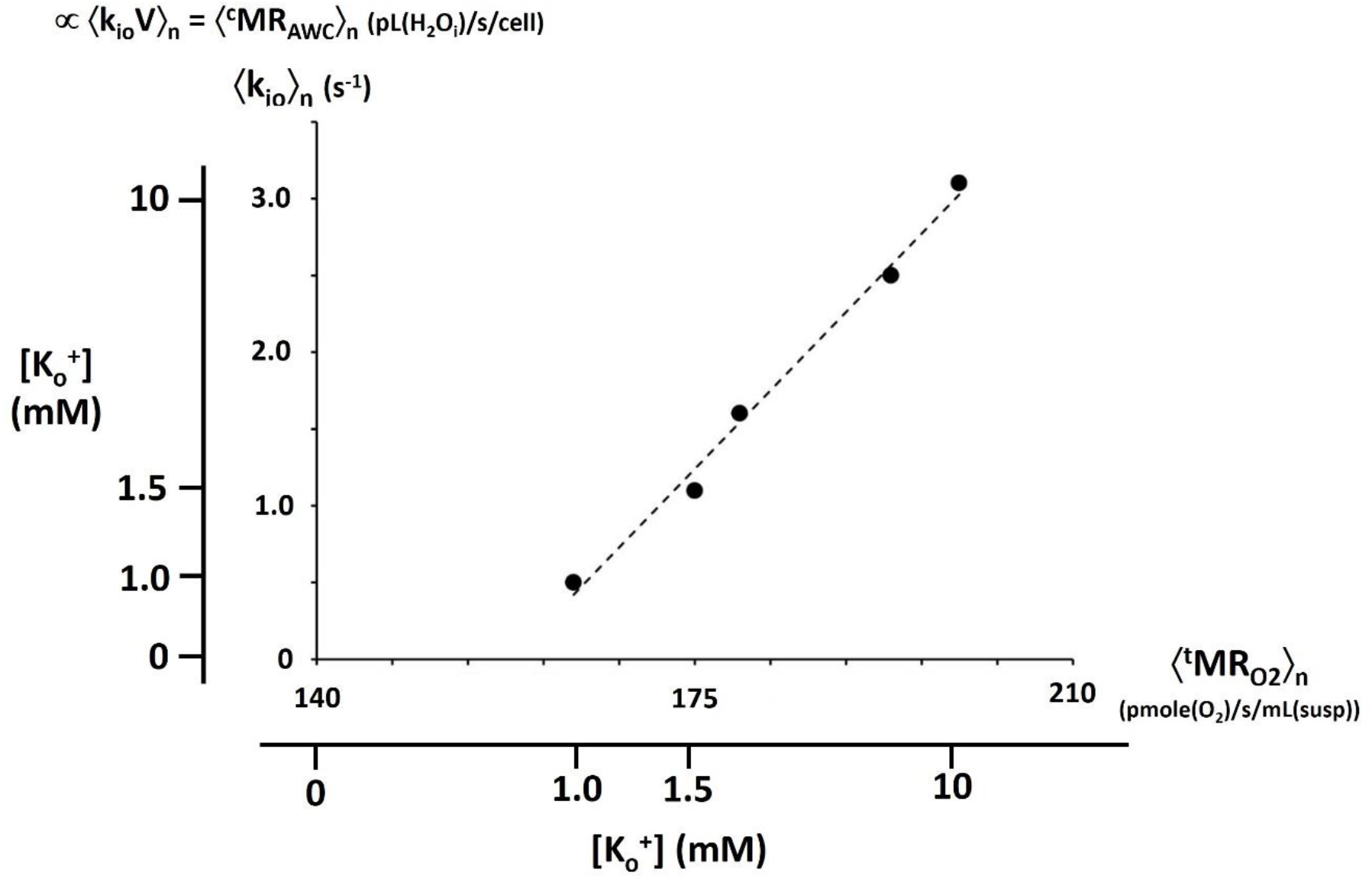
A correlation of k_io_ with ^t^MR_O2_. Independent [K_o_^+^] titrations of paramagnetic agent-superfused [SS-NMR] studies of organotypic, cultured [spiking] rat somatosensory cortex [ordinate] and direct studies of isolated rat brain synaptosome suspensions [abscissa] allow correlation of the population-averaged ⟨k_io_⟩_n_ and ⟨^t^MR_O2_⟩_n_ quantities, respectively. The outer [K_o_^+^] scales are non-linearly related to the linear, inner ⟨k_io_⟩_n_ and ⟨^t^MR_O2_⟩_n_ scales due to their Michaelis-Menten relationships. The correlation of ⟨k_io_⟩_n_ and ⟨^t^MR_O2_⟩_n_ is very strong. The SS-NMR studies also indicate the mean cell volume ⟨V⟩_n_ is rather constant. Thus, ⟨k_io_V⟩_n_ = ⟨^c^MR_AWC_⟩_n_ (pL(H_2_O_i_)/s/cell) correlates with ⟨^t^MR_O2_⟩_n_ (pmole(O_2_)/s/mL(suspension)). (This is a combination of Figures 3C and 3E of reference (31), where details are provided. The points are synaptosome measurements: the dashed line is the Michaelis-Menten fitting (K_m_ = 4.5 mM) of the cortical culture measurements.)

In Figure 2, AWC participation of Fig. 1 enzymes is exemplified by SGLT (both a type III and a type IV transporter) and by KCC4 (a type II and type I enzyme). These transporters share substrates with NKA; Na_o_^+^ for SGLT and K_i_^+^ for KCC4. The SGLT and KCC4 reactions are thereby coupled with the NKA reaction. As individual proteins, I and IV respectively are catalyzing *unidirectional* water effluxes and influxes. Working in concert with other transporters, and orchestrated by NKA activity however, they produce overall cellular homeostatic water *bidirectional* exchange. The cellular AWC fluxes are given in **Equation (2)**. This is a “systems” aspect of the cell. Though none of the individual process metabolic

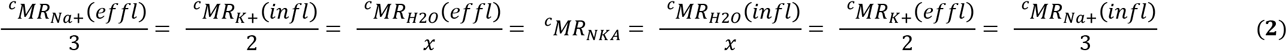

rates (^c^MR’s) are equal to ^c^MR_NKA_, they rise or fall with the latter. For example, ^c^MR_K+_(effl) = 2^c^MR_NKA_ and ^c^MR_H2O_(effl) = x^c^MR_NKA_ = [H_2_O_i_]k_io_V. Thus, ^c^MR_H2O_(effl) = ^c^MR_H2O_(infl) = x^c^MR_NKA_. NKA is the driver.

### Molecular Dynamics (MD)

*In silico* simulations of Fig. 1 transporter mechanisms using MD (29,30,32-37) can be very informative. This is particularly so for studies of an SGLT enzyme: the bacterial transporter vSGLT forms a hydrophilic membrane-spanning channel (32-36). Generally, the enzyme exhibits spasmodic (tens of ns) bursts, some featuring *net* water unidirectional influx and some featuring bidirectional water *exchange* (33-35). That is, a given transporter can indeed facilitate the net flux (category B) and the exchange (category A) processes at different times (32).

For a period after sodium ion release in the vSGLT sugar-bound state, water molecules pass inward and outward, past the galactose molecule (a glucose stereoisomer) in the channel, in almost equal, small numbers (∼one molecule/ns) - homeostatic exchange. The sugar is bound to its site midway through the channel, while a Na^+^ binding site is near the cytoplasmic mouth. In the ∼100 ns after galactose is released, it proceeds through the channel into the cytoplasm, and pushes the channel-filling water molecules before it <u>into</u> the cell - net influx - in a quantity (33,35) generally consistent with the Fig. 1 SGLT water stoichiometries. Both such periods (exchange and net flux) are only transient (34). However, both seem triggered by Na^+^ release.

Since the common role of the Fig. 1 enzymes is to facilitate passage of hydrophilic substrates across the hydrophobic bilayer membrane, it is likely they all have somewhat similar hydrophilic pathways. The structure of the NKCC1 transporter channel has been determined (38). We surmise there are many more water co-transporting membrane enzymes, possibly including Na^+^,K^+^-ATPase (NKA) itself, as yet unstudied for this aspect. The NKA channel structure has been elucidated (39). Transient exchange periods simulated for AQP-1 (37) are very similar to those for vSGLT (34). It is likely each of the Fig. 1 transporters can accomplish what AQPs accomplish, but in addition the conduct of specific metabolites across the membrane. Water molecules will find their ways through any open hydrophilic pathway (“pore”).

The aforementioned vSGLT MD simulations also suggest possible molecular mechanisms for SGLT/NKA *kinetic* coupling. As stated, large tranches of net water molecule influx (category B) generally occur only upon galactose release from its site near the middle of the hydrophilic channel. But, the probability for this, and for exchange (category A), can depend on whether a Na^+^ ion is in a site: the Na^+^ can block the sugar (and apparently *net* water) ingress into the cell (35,36). More often than not, Na^+^ release occurs before sugar release (32). At least 80 ns after the Na^+^ release (35,36), galactose release can occur – unless the Na^+^ ion has re-bound or has been replaced by another Na^+^ ion. While galactose is still present, the duty cycle of the Na^+^ site remaining empty depends on the intracellular sodium concentration [Na_i_^+^]: this Na^+^ can come from only inside the cell. With a normal ^c^MR_NKA_, [Na_i_^+^] is small (see below). However, [Na_i_^+^] increases as ^c^MR_NKA_ decreases (40). Thus, the probability of the Na^+^ site being re-occupied within 80 ns increases as ^c^MR_NKA_ decreases. As a consequence, the sugar-release frequency decreases as ^c^MR_NKA_ decreases. In this way, MR_SGLT_ and MR_NKA_ are positively correlated kinetically. (This pair-wise SGLT/NKA consideration assumes all other Figs. 1,2 enzyme activities remain unchanged.) Irrespective of the details, the sharing of any substrate with NKA should facilitate kinetic coupling. The correlation mechanisms of other Fig. 1 water co-transporter activities with MR_NKA_ must bear some general similarity to this. Pore openings and closings are metabolically controlled.

### A Futile Cycle?

When they were first realized, the Fig. 2 Na^+^ and K^+^ exchange processes may have been briefly viewed as “futile cycles.” This term is used to characterize a set of coupled biochemical reactions that consume ATP with no apparent benefit (41). Obviously, it is generally thought biological evolution discards futile cycles as “energy wasting.” However, it was quickly understood the trans-membrane ion concentration gradients enjoyed by Na^+^ and K^+^ serve to partially store chemical (Gibbs) free energy released by NKA-catalyzed ATP hydrolysis, ΔG_ATP_ = **-** 59 kJ/mole (42,43). As we will see below, ΔG_Na_(infl) is typically near – 14.5 kJ/mole, for Na^+^ influx, and ΔG_K_(effl) near 0 kJ/mole, for K^+^ efflux. Thus, though these cycles are *homeostatic*, they are not in *equilibrium*, but in *steady-state*. Energy is required to maintain them. Subsequently, the free energy thus stored in the [Na^+^] gradient is used to facilitate cellular uptake of many crucial molecules (Fig. 1), and these processes help complete the Na^+^ cycle. The free energy in the [K^+^] gradient is small only because of the cell’s membrane potential, which the homeostatic K^+^ efflux largely produces and maintains (see below).

The “pump and leak mechanism” term used for water (19,27) implies waste. However, a question naturally arises. Might the existence of active trans-membrane water cycling (Fig. 2) reveal an analogous metabolic energy storage function? Is AWC an actual example of a futile cycle, or does it have some evolutionary advantage?

## THEORY

### Reaction Free Energy Changes (ΔG)

The issue of AWC futility is a chemical thermodynamic question. Thus, we must consider ΔG for each permeant as it crosses the membrane, for example, into the cell; ΔG_per_(infl), for *permeant* (per) influx (infl). For transport reactions (as in Fig. 1), the standard free energy change (ΔG^0^) is zero. There are potential chemogenic, electrogenic, and barogenic ΔG contributions. These are, respectively, the three terms on the right-hand-sides of **Equations (3)**; where i and o are, respectively, the inside and outside compartmental indices, [per] is

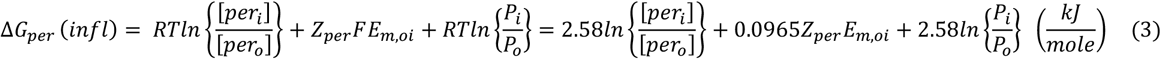

the compartmental permeant concentration, Z_per_ the (signed) permeant particle electrical charge, F is the Faraday constant (0.0965 kJ/(mV•mole) at physiological temperature, 310 K), E_m,oi_ is the (signed) trans-membrane electrical potential (in minus out) in mV, P_i_/P_o_ is the intracellular/extracellular hydraulic pressure ratio (Figs. 1,2) ((44), p. 35), and RT = 2.58 kJ/mole at 310 K. For H_2_O as permeant, Z_H2O_ = 0, and the second term makes no contribution.

Equations (3) can be written alternatively with permeant *chemical potential* changes; Δμ_per_, where μ_per_≡ (∂G/∂n_per_)_T,P,n(≠ per)_ ((44), p. 20). However, the ΔG notation is more inclusive, and more common in this field (42,43).

To obtain the free energy change for any Fig. 1 *reaction*, ΔG_reac_(infl), one must sum over all permeants, **Equation (4)**; where s_per_ is the permeant stoichiometric coefficient (Fig. 1). It is obvious from Eqs. (3,4) the chemogenic

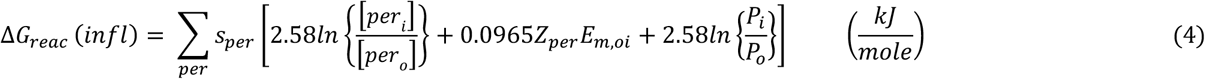

terms require realistic compartmental permeant concentrations. Strictly, thermodynamic activity (a = the concentration•activity coefficient product) values should be used. Below, and in **A2**, we justify the first-order approximation that solute and solvent activity coefficients are each unity.

### Concentration Scales

In this paper, a concentration is indicated with square brackets, *e*.*g*., [permeant]. Sometimes, the molarity scale, with unit M = moles (permeant)/L(compartment) (activity coefficient y_per_ ((45), pp. 26,27)) is used. However, first-order evaluation of chemogenic terms in Eqs. (3) and (4) consider only the entropy of mixing. This requires concentration scales with mole ratio- or mole fraction (X)-like quality. Molality, with unit m = moles (permeant)/kg(water) (activity coefficient γ_per_ ((45), pp. 26,27)), is one such scale. Biological aqueous solutions are sufficiently dilute in each solute that its m value is well-approximated by its M value; the *infinite dilution* assumption. The use of an entropy-centric concentration scale here is tantamount to assuming solution “ideality;” there are no specific solute/solute interactions. We will see below this is clearly not the case. The “volume molality” scale, _V_m = moles(permeant)/L(solvent) ((44), p.37) will also be considered. (Using m or _v_m concentration values is tantamount to assuming the small molecule metabolites may sample the entire compartmental space or only aqueous spaces, respectively, (*i*.*e*., the water-excluding macromolecules are more or less mobile or immobile (14).) Since all the different neutral and ionic solutes (thus, osmolytes plural) need be counted, we will further use the OsMolarity (unit, OsM), or Osmolality (unit, Osm) concentration scales.

Since we also consider the solvent water as a permeant (Fig. 1), we must express its concentration. For understanding water thermodynamic activity – its “*escaping tendency*” - the dimensionless X (or a related) scale is best ((44), p. 56; (45), p.27; (46)). Thus, for water, X_H2O_ = (moles(water))/(moles(osmolytes + water)), activity coefficient f_H2O_ ((45), pp. 26,27)), is used.

One will further see milli-scales: mOsM, mOsm, and mOsX.

### Chemogenic Contributions

#### Tissue Osmolyte Compartmental Concentrations

For *in vivo* conditions, accurate determination of biological compartmental solute concentrations is difficult, particularly for intracellular metabolites. Recent metabolomic and proteomic advances (for example, using calibrated liquid chromatography/mass spectrometry techniques) have given realistic accountings. From such a study of cultured murine kidney epithelial cells, 101 different intracellular small molecule metabolites, and their concentrations, have been itemized (47). We compile an inventory of intra- and extra-cellular solute concentrations from a number of related studies (47-62) in **Table 1**. The inorganic ion (Na^+^, K^+^, H_3_O^+^, Ca^2+^, Mg^2+^, Cl^**-**^, HCO_3_^**-**^) concentrations are conventional. The units are mOsM (thus, mOsm for osmolytes). These should be considered representative of interstitial and intracellular compartment osmolytes. Of course, they are not the values for any actual cell. Concentrations surely differ from cell to cell, tissue to tissue, and in different biological conditions. Since we are calculating general thermodynamic trends, the Table 1 values suffice.

**Table 1.** Representative Osmolyte Concentrations (mOsM)^a-d^.

| Table 1. Representative Osmolyte Concentrations (mOsM)<sup>a-d</sup> |  |  |
| --- | --- | --- |
|  | interstitial | intracellular |
| Na<sup>+</sup> | 135.0 | 15.0 |
| K<sup>+</sup> | 5.0 | 150.0 |
| H<sub>3</sub>O<sup>+</sup> | 0.00005 | 0.0001 |
| Ca<sup>2+</sup> | 1.5 | 0.0001 [free]; 0.01 [total] |
| Mg<sup>2+</sup> | 0.5 | 0.5 [free]; 10 [total] |
| other cations | 1.5 | 3.5 |
| Cl<sup>-</sup> | 105.0 | 15.0 |
| HCO<sub>3</sub><sup>-</sup> | 30.0 | 10.0 |
| small molecule metabolite anions | 3.0 | 135.0<sup>a,b</sup> |
| protein anions | 7.5<sup>a</sup> | 9.5<sup>a,c,d</sup> |
| small molecule metabolites |  | 40.0<sup>a,b</sup> |
| RNA |  | 0.5<sup>d</sup> |
| lipids |  | 0.05<sup>d</sup> |
| DNA |  | 0.000005<sup>d</sup> |
| total osmolytes | 289 | 379 |
| H<sub>2</sub>O mole fraction (OsX<sub>w</sub>) | 0.995 | 0.993 |
| H<sub>2</sub>O millimolarity (mM) | 45,000<sup>e</sup> | 42,000<sup>f</sup> |
| f<sub>w</sub> | f<sub>w,o</sub> = 45/55 = 0.82 | f<sub>w,i</sub> = 42/55 = 0.76 |
| osmotic stoichiometry (H<sub>2</sub>O/osmolyte) | 45/0.289 = ~156 | 42/0.379 = ~111 |
| References (47-62). <sup>a</sup>ref. (48), <sup>b</sup>ref. (47), <sup>c</sup>ref. (49), <sup>d</sup>ref. (50), <sup>e</sup>ref. (62), <sup>f</sup>ref. (51). |  |  |

We see the intracellular RNA, lipid assembly, and DNA mOsM concentrations are tiny. Though these occupy considerable intracellular volume (see below), because of their very large macromolecular masses they make negligible osmolal contributions. Since there are fewer (but non-zero: see below) extracellular macromolecules, we ignore their osmolal contributions here.

Interestingly, we see the total osmolyte concentrations are ∼0.3 OsM for the interstitial space and ∼0.4 OsM for the intracellular space. It is widely assumed intracellular osmolarity is near 0.3 (52), and thus similar to that in the extracellular space. The implications of this trans-membrane osmolarity gradient are explored in the section on intracellular pressure (P_i_) below.

#### Tissue Water Compartmental Concentrations

Experimentally determining biological compartmental water contents is even harder than for solutes (2). For many years, it has been stated, usually anecdotally, that cell water content is 70% (v/v) [or (w/w), assuming unit density, d] (53). Vinnakota and Bassingthwaighte presented a comprehensive review and tabulation of rigorous, quantitative determinations of myocardial tissue compartmental densities, volumes, volume fractions, and mass fractions (51). These derive from direct *ex vivo* mass analyses and *in vivo* tracer studies, and yield the cell water content value 75 % (w/w). A more recent optical determination yields 85 % (v/v) for cultured mammalian cell controls (54).

If we use the 75 % value (51) and take d = 1.1 g/mL, we obtain [H_2_O_i_] = 42 M; *i*.*e*., 42 moles(H_2_O)/L(cell). This is the most parsimonious interpretation of the data, and includes water in the cytoplasm, mitochondria, and sarcoplasmic reticula. The magnitude is considerably reduced from the value for pure water, 55 M. The intracellular water volume fraction, f_W,i_, estimated from this is 42/55 = 0.76 (Table 1). This is in good agreement with quantitative MRI *in vivo* measures (55). (Note: f_W_ is different from f_H2O_.) Rand states, without attribution, [H_2_O_i_] = 54 M (56).

From tracer (^15^OH_2_) studies (reviewed in (51)) and NMR (^1^H_2_O) studies (reviewed in (2)), essentially all intracellular water is effectively “well-mixed,” at homeostasis. There has been much consideration of “bound” water in tissue (reviewed in (25,57-59)). There is no doubt water near macromolecule and membrane surfaces is thermodynamically different from bulk water – it has at least lower entropy (see below) – but kinetically (tumbling- and diffusion-wise) it remains in very rapid communication with all the water in the cell. The fraction of water molecules in “buried” intra-macromolecular sites inside cells is miniscule. Though the latter have significant NMR consequences (reviewed in (25)), even these water molecules have labile access to the cytoplasm (60).

Because of its small volume fraction (typically, 0.2) the extracellular compartmental water content in parenchymal tissue has been even more difficult to determine. Vinnakota and Bassingthwaighte report myocardial *interstitial* water content 92 % (w/w) and d = 1.0 g/mL, but this has a significantly greater uncertainty (51) and seems excessively large. The water content of cartilage can be more directly determined, and this might be a good extracellular model: it is an essentially acellular matrix of collagen fibrils (61). Lu and Mow imply a cartilage water content value of 78 % to 85 % (61). Maroudas and co-workers carefully measured the value 81 % for unloaded human femoral head cartilage (62). Again assuming unit density, this yields [H_2_O_o_] = 45 M, also significantly reduced from 55 M, though not as much as [H_2_O_i_]. The corresponding f_W,o_ is 45/55 = 0.82 (Table 1).

In Table 1, we give intra- and extracellular “osmotic stoichiometry” values; the overall ratio of water molecules to osmolyte particles in the compartment. We see the intra- and extracellular values are near 100 and 150 water molecules per osmolyte, respectively. The fact that in Fig. 1 transporter osmotic stoichiometries for the reactants are either above or below these values is consistent with the water stoichiometry being principally determined by the hydrophilic channel volumes (35). The molecular action of any particular transporter has nothing to do with the overall “osmotic stoichiometry.” It is only the *net* water transported by all influxers and effluxers in the cell that must be zero for homeostasis. However, even if values of 100 or 150 did obtain, this would be irrelevant for the calculations below.

The large macromolecular volumes are the reason the 42 M intracellular H_2_O concentration (Table 1) used below is so much less than that of pure water (55 M). These large molecules exclude considerable water (Fig. 8 of (15)). Obviously, there are also water-displacing extracellular macromolecules: 45 M (Table 1) is also smaller than 55 M.

However, the molar concentration scale is useful mainly for molecular counting (1 mole = 6 x 10^23^ molecules). As justified above, we use the dimensionless mole fraction concentration scale for water (X_H2O_). (This differs from the tissue compartmental water mole fractions (“populations”) p_i_ and p_o_ (important in SS-NMR).) For most practical aqueous solutions, including those in biological compartments, however, X_H2O_ is always very nearly unity. “In dilute solutions of electrolytes, the activities, and even the activity coefficients (f_H2O_), of the *solvent* are little different from unity” ((63), p.12). Very important consequences of this are considered below.

One kg of water is (1000/18.0154) 55.508 moles H_2_O. Dividing this by (55.508 + the sum of osmolyte concentrations) in each of the cytosolic and interstitial compartments yields corresponding water mole fractions OsX_H2O,i_ = 0.993 and OsX_H2O,o_ = 0.995 (Table 1). OsX_H2O,i_ is only 0.2% smaller than OsX_H2O,o_, and each value is nearly that for pure water. Given this situation, following Pitzer ((63), p.12), it is not unreasonable to assume equal activity coefficients (f_H2O,i_ = f_H2O,o_ = 1) and thus use these mole fraction values in the Eqs. (3) and (4) terms for water as permeant. Inserting {OsX_H2O,i_/OsX_H2O,o_} = 0.998 yields ΔG_H2O_(infl) = **-** 0.00517 kJ/mole for the *chemogenic* contribution.

Therefore, unlike the other Table 1 species, water is almost (but not exactly) in trans-membrane chemical equilibrium. (In this paper, “*chemical*” equilibrium is defined as: [per_i_] = [per_o_].) There is (only) ∼5 J/mole favoring cellular water influx. Strictly, this chemogenic ΔG_H2O_(infl) value might be considered not significantly different from zero (chemical equilibrium). However, analyses of surrogate solutions in **A2** give water activity values: a_H2O,i_ = 0.994 and a_H2O,o_ = 0.995. These correspond to chemogenic ΔG_H2O_(infl) = **-** 0.003 kJ/mole. Henceforth, we will use – 3 J/mole. As we will see, this has no bearing on the important metabolic consequences detailed below. For co-transported water, the barogenic contribution will be much greater than the chemogenic term.

It is probably unreasonable that chemogenic ΔG_H2O_(infl) be positive. First-order osmotic considerations more strongly favor water influx. However, the first-order is inadequate, see below.

### Electrogenic Contributions

We make the common approximation that K^+^ *efflux* permeabillity dominates over Na^+^ and Cl^**-**^ permeabilities (in the P_f_ sense) in cellular homeostasis. Thus, we use a version of the Nernst **Equation (5)** ((44), p. 113) to calculate E_m,oi_,

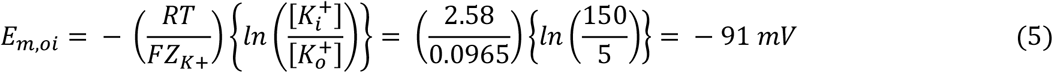

where: RT = 2.58 kJ/mole and the Faraday constant F = 0.0965 kJ/mV•mole (both at 310 K) and insert the Table 1 [K^+^] values. Z must be the value for the potential-producing ion, Z_K+_ = 1. It is important to note the actual number of *excess* K_o_^+^ ions (and an equal number of uncompensated intracellular anions) is insufficient to significantly alter the Table 1 [K^+^] values. An excess of only 0.02 mM intracellular anionic charges (0.014% = ((100×0.02)/145); Table 1) produces E_m,oi_ ≈ - 100 mV for a small (1 fL) spherical cell ((53), p. 198.)

### Barogenic Contributions

Intracellular pressure (P_i_) can be counteracted with an externally applied (macroscopically uniform) mechanical pressure, in principle ((44), p.32), and elegantly transduced into an isotropic, hydraulic (mechanical) pressure with cellular insertion of a pressure-sensitive “osmotic” microelectrode (64,65). The trans-membrane pressure difference (P_i_ – P_o_) force is oriented roughly vectorially radial to the cell surface. Experimentally, most mammalian cells have (P_i_ - P_o_) values under 0.1 atm (27). However, an osmotic microelectrode study of perfused *ex vivo* murine lens (featuring relatively simple lens cells) found (P_i_ - P_o_) values up to 0.5 atm (66). Most P_o_ values are near 1 atm: even “high” tumor interstitial P_o_ is only 1.02 atm. (67). For water, the standard state (f_H2O_ truly 1) pressure, P^0^, is 1.0 atm.

## RESULTS

### Free Energy Calculations

#### *Permeant* Influx Free Energy Changes, Eq. (3)

In **Table 2**, we list many potential non-metabolite permeant (per) species, and calculate chemogenic, electrogenic, and barogenic ΔG(infl) contributions (kJ/mole) for their influx. For these calculations, we use the Table 1 water OsX_W_ and solute mOsm concentrations for the chemogenic terms. We calculate the barogenic contributions for P_i_/P_o_ ratios of 1.00 (blue) and 1.05 (red).

**Table 2.**
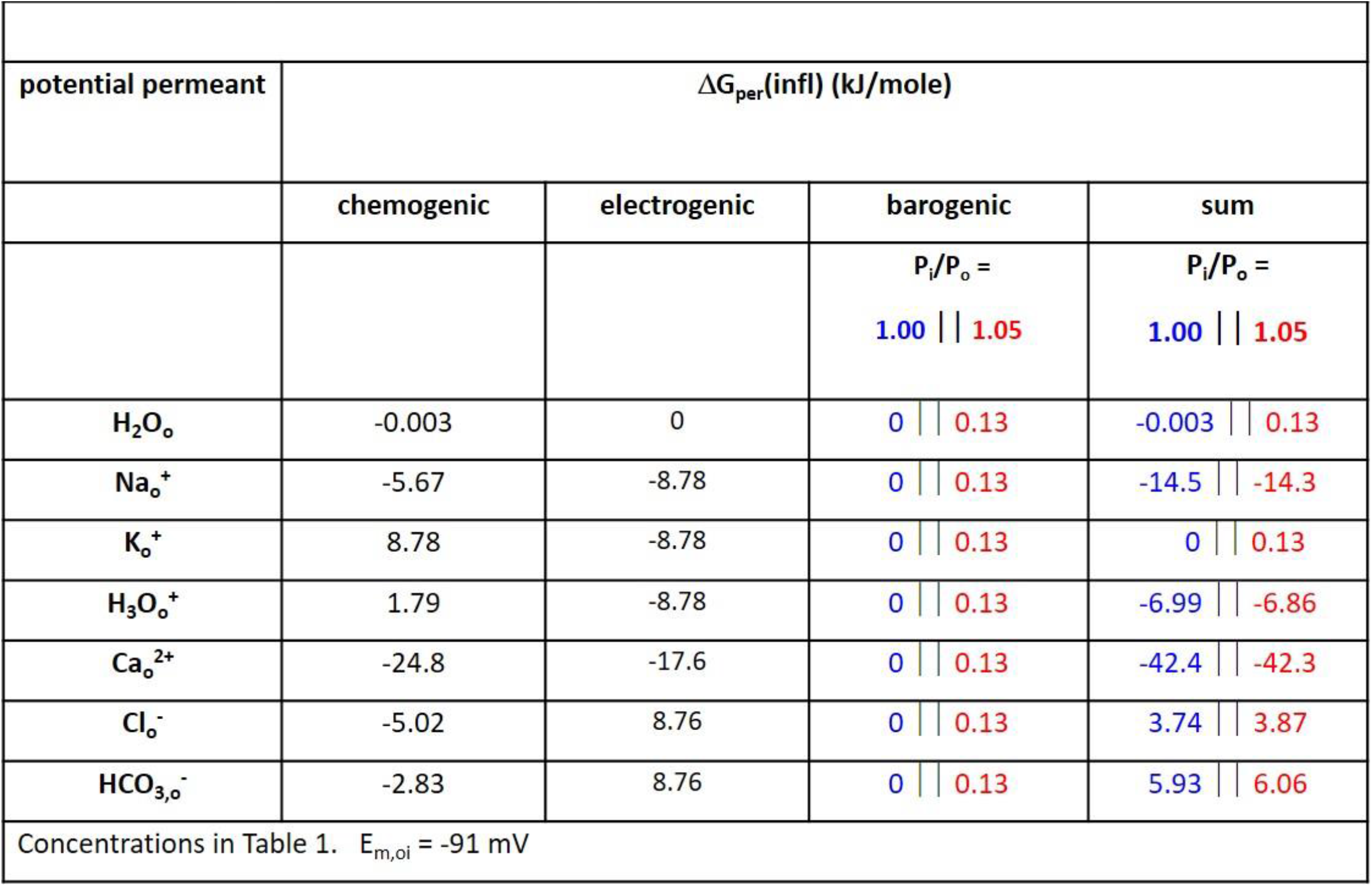
Free Energy Change for Influx, Eq. (3).

It is clear that, compared with those of the other permeants, the chemogenic and electrogenic contributions for individual H_2_O molecule transport are negligible. We see the slight chemogenic tendency for water molecule influx when P_i_ = P_o_is reversed when P_i_ ≥ 1.05 P_o_. In fact, for ΔG_H2O_(infl) = **-** 0.003 kJ/mole, any P_i_ > 1.001 P_o_ would result in water efflux if there were no other considerations.

Let us consider two other potential permeants, Na_o_^+^ and K_o_^+^ as examples. The electrogenic terms for their *influxes* are identical, **-** 8.78 kJ/mole. Electrostatically, influx of each is favored. However, while the chemogenic Na_o_^+^ contribution (**-** 5.67 kJ/mole) also favors influx, the chemogenic K_o_^+^ term (8.78 kJ/mole) opposes influx. Thus, the *electrochemical* potential gradient for Na_o_^+^ influx is very favorable, – 14.5 kJ/mole (**-** 5.67 – 8.78), but zero for K_o_^+^ influx (or K_i_^+^ efflux) when P_i_ = P_o_. The barogenic contributions for Na_o_^+^ and K_o_^+^ are identical, but this has very small effect for only Na_o_^+^. However, compared with Na_o_^+^, and indeed with all the other Table 2 permeants except K_o_^+^, the barogenic contribution for H_2_O is *proportionally* much greater than the chemo- and electrogenic terms. (For example, a P_i_ of 1.05 atm causes a more than 40-fold increase in ΔG_H2O_, but only a 1% change in ΔG_Na_.) This has very important consequences.

#### Transport *Reaction* Free Energy Changes, Eq. (4)

We use Eq. (4) to calculate the total ΔG_transporter_(infl) for each of the Fig. 1 transporter reactions - utilizing the stoichiometries seen there. The “metabolite” substrates (glucose, glutamate^**-**^, GABA, and lactate^**-**^) are omitted because we study below how their transport is affected by the thermodynamic contributions of the other “ancillary” (support) substrates.

##### Influx Reactions

Reactions that normally run in the *influx* direction (IV, III, or both, in Figs. 1,2) are presented in **Table 3a** (E_m,oi_ is fixed at – 91 mV). Each is favorable (negative ΔG) when P_i_ = P_o_ (blue). However, they each become less favorable as P_i_/P_o_ increases, and all become unfavorable (positive ΔG) when P_i_ has reached 1.05 P_o_ (red). This is due to P_i_ opposition to water co-influx (the factors that are - 0.003 when P_i_ = P_o_ become positive when P_i_ exceeds 1.001 P_o_). Because of the large water stoichiometries, the barogenic contribution can have a significant influence on secondary active water co-transporter thermodynamics. This is due to essentially only the water co-transport. For the other permeants, the blue/red values generally have very similar magnitudes.

**Table 3a.**
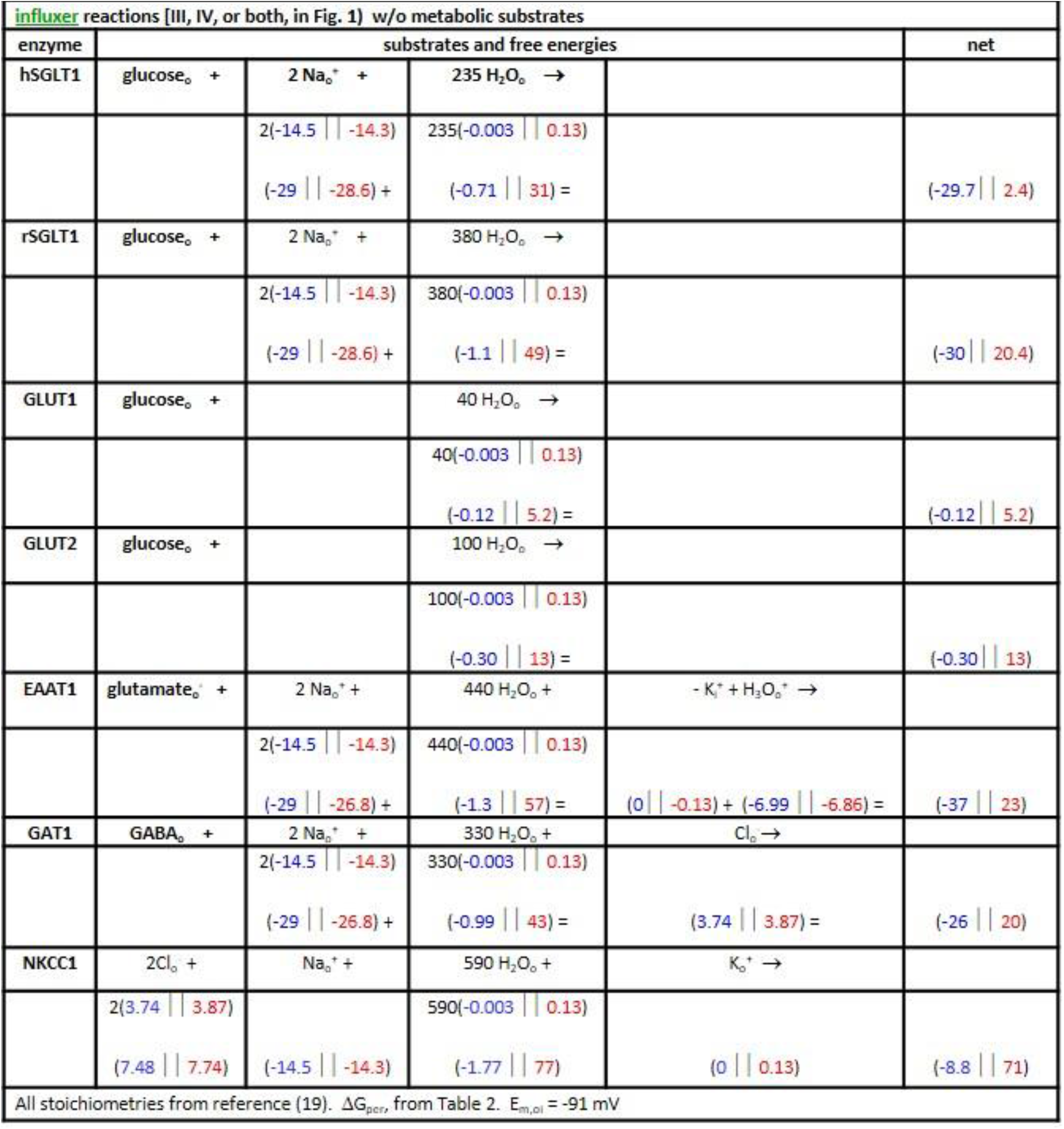
Transport Reaction Free Energy Changes, ΔG(infl) (kJ/mol); Eq. (4) (P_i_/P_o_ = 1.00 | | 1.05)

**Table 3b.** Transport Reaction Free Energy Changes, ΔG(effl) (kJ/mol), Efflux Version of Eq. (4) (P_i_/P_o_ = 1.00 | | 1.05)

| KCC4 | $K_i^+ +$ | $Cl_i^- +$ | 500. $H_2O_i \rightarrow$ | | net |
| --- | --- | --- | --- | --- | --- |
| | | | 500(0.003 \| $-0.13$) | | |
| | (0 \| $-0.13$) + | (-3.74 \| $-3.87$) + | (1.5 \| $-65$) = | | (-2.2 \| $-69$) |
| MCT1 | $\text{lactate}_i^- +$ | $H_3O_i^+ +$ | 500 $H_2O_i \rightarrow$ | | |
| | | | 500(0.003 \| $-0.13$) | | |
| | | (6.99 \| $6.86$) + | (1.5 \| $-65$) = | | (8.5 \| $-58$) |
| NaDC1 | $\text{dicarboxylate}_i^{2-} +$ | | 175 $H_2O_i +$ | 2 $Na_i^+ \rightarrow$ | |
| | | | 175(0.003 \| $-0.13$) | 2(14.5 \| $14.3$) | |
| | | | (0.53 \| $-23$) + | (29 \| $29$) = | (30 \| $6.0$) |
| All stoichiometries from reference (19). $-\Delta G_{\text{per}}(\text{infl})$ from Table 2. $E_{m,oi} = -91$ mV | | | | | |

##### Glucose Uptake at Fixed [glucose_i_]

For the rSGLT1 influx reaction as an example: ΔG = **-** 30 kJ/mole when P_i_ = P_o_, but 20.4 kJ/mole when P_i_ = 1.05 P_o_. (P_i_ = 1.03 P_o_ is sufficient to make influx unfavorable.) To illustrate the consequences of this on glucose uptake, we fix the intracellular glucose concentration, [glc_i_], at 2 μM. (The intracellular enzyme that phosphorylates glucose, hexokinase, has a glucose_o_ K_m_ value reported as 1.7 μM (20). A principle of central carbon (glucose-related) metabolism is that any intracellular metabolite steady-state concentration should be near the K_m_ value of the enzyme that consumes it (47).) Thus, we write Eq. (4) explicitly for the 2Na^+^/glucose co-transporter **rSGLT1** influx free energy change, ΔG_rSGLT1_(infl), in **Equation (6)**. The first term represents the glucose barochemical contribution

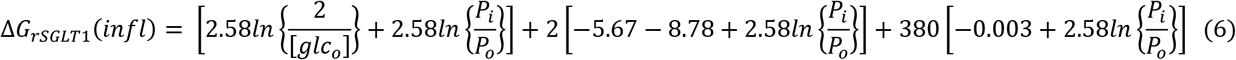

with [glc_i_] fixed at 2 μM. The second and third terms are due to, respectively, the Na^+^ electro- and barochemical and the H_2_O barochemical potential gradients. The former favors influx, while the latter opposes it.

If we fix P_i_ = 1.05 P_o_, and neglect the water co-transport, ΔG becomes positive (*i*.*e*., influx ceases) only when the extracellular glucose concentration [glc_o_] decreases below 0.33 nM. On the other hand, if we keep P_i_ = 1.05 P_o_ but retain the water co-transport, the reaction becomes unfavorable when [glc_o_] decreases below 2.4 mM – three orders of magnitude *greater than* [glc_i_]. The large electrochemical ΔG_Na+_ favoring influx (**-** 28.6 kJ/mole) is counteracted by the even larger barochemical ΔG_H2O_(infl) favoring efflux (49 kJ/mole). A number of significant consequences of this will be considered in the Discussion section.

To partially generalize these considerations, **Figure 4** shows a 3D plot of Eq. (6) for the rSGLT1 reaction. For this, the [Na_o_^+^] and [Na_i_^+^] values were fixed at those in Table 1, [glc_i_] at 2 μM, and E_m,oi_ at – 91 mV. The vertical axis plots the rSGLT1 influx reaction ΔG_rSGLT1_(infl), while the (logarithmic) oblique axes increment [glc_o_] and P_i_/P_o_. The free energy surface is colored green when influx is possible (ΔG_rSGLT1_(infl) < 0), and red when impossible (ΔG_rSGLT1_(infl) > 0). The intersection of the ΔG_rSGLT1_(infl) surface with the horizontal ΔG_rSGLT1_(infl) = 0 plane defines the trajectory of the (P_i_/P_o_)-dependence of the minimum extracellular glucose concentration required for influx. While the exponential nature of the (P_i_/P_o_)-dependence is hardly noticeable, that of [glc_o_] is very evident.

**Figure 4.**
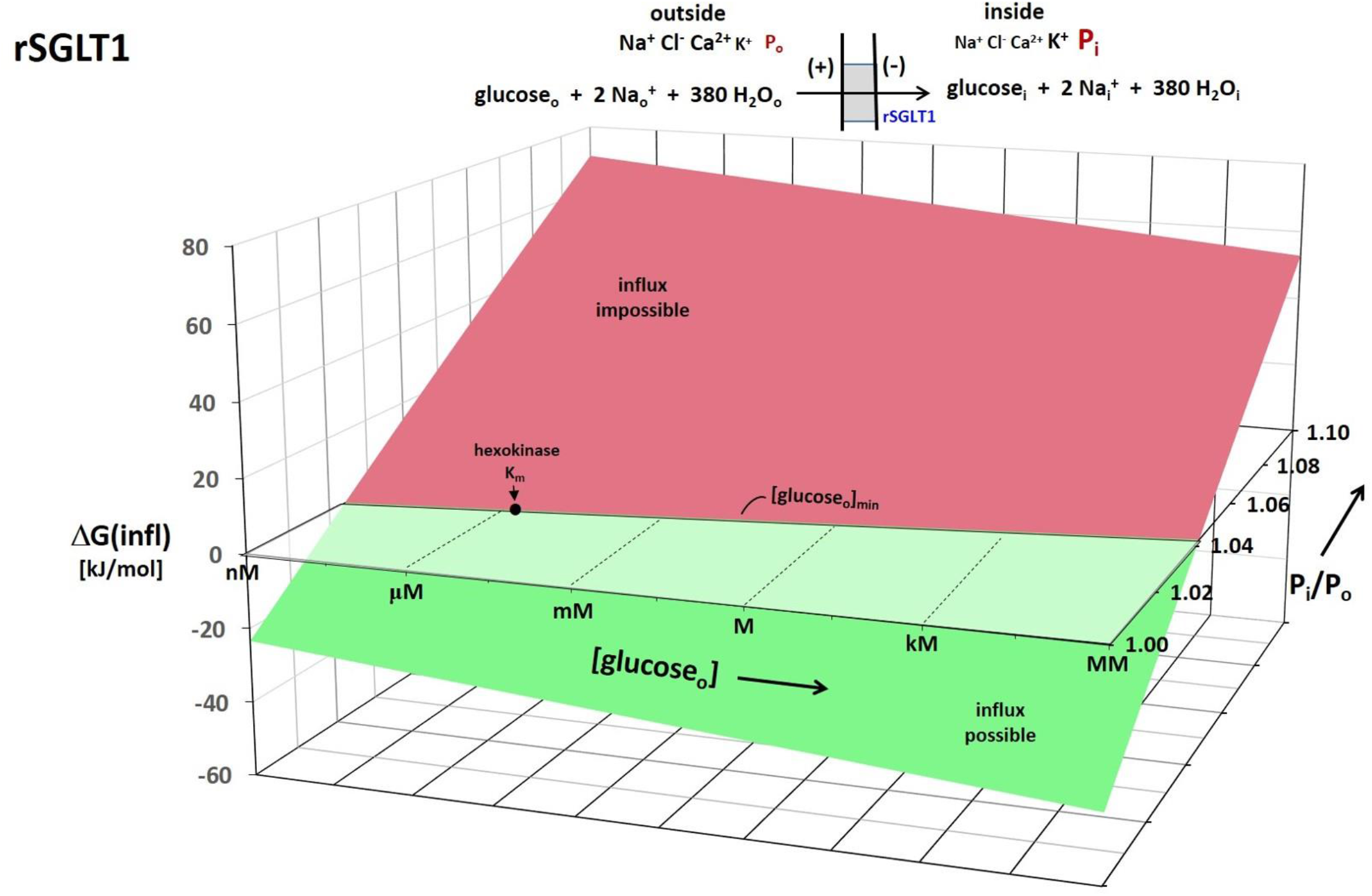
A 3D plot for the rSGLT1 reaction. The vertical axis measures the Gibbs free energy change (ΔG) for the influx direction shown, and calculated with Eq. (6). The logarithmic oblique axes plot the extracellular glucose concentration, [glucose_o_], and the intracellular/extracellular hydraulic pressure ratio (P_i_/P_o_) over the experimentally measured range. For this calculation, the Table 1 concentrations and the Tables 2 and 3a free energy terms were used (chemogenic ΔG_H2O_(infl) = **-** 3 J/mole). The intracellular glucose concentration, [glucose_i_], the membrane potential, E_m,oi_, and P_o_ were held fixed at 2 μM, **-** 91 mV, and 1 atm, respectively (T = 310 K). The surface is colored green when influx is thermodynamically possible and red when it is impossible. Thus, the intersection of the ΔG surface with the ΔG = 0 plane traces the trajectory of the (P_i_/P_o_)-dependence of the *minimum*, [glucose_i_]_min_, value. The value of the intracellular hexokinase K_m_ (1.7 μM) for glucose_i_ is indicated.

To make these effects clearer, **Figure 5** shows the 2D plot of the Fig. 4 ΔG_rSGLT1_(infl) = 0 plane. With H_2_O co-influx, increasing P_i_/P_o_ strongly increases [glc_o_]_min_. For P_i_ = 1.04 P_o_, [glc_o_]_min_ is already ∼10 μM (*greater than* [glc_i_] here) for rSGLT1. The location of the hexokinase glucose_i_ K_m_ is shown as a horizontal dashed line. Of course, the position of the surface depends also on the E_m,oi_, [glc_i_], and [Na_i_^+^] values, among other quantities. For example, making ΔG_Na_(infl) less negative (say, by increasing [Na_i_^+^]) would increase [glc_o_]_min_, all other factors being equal – a given P /P, for example.

**Figure 5.**
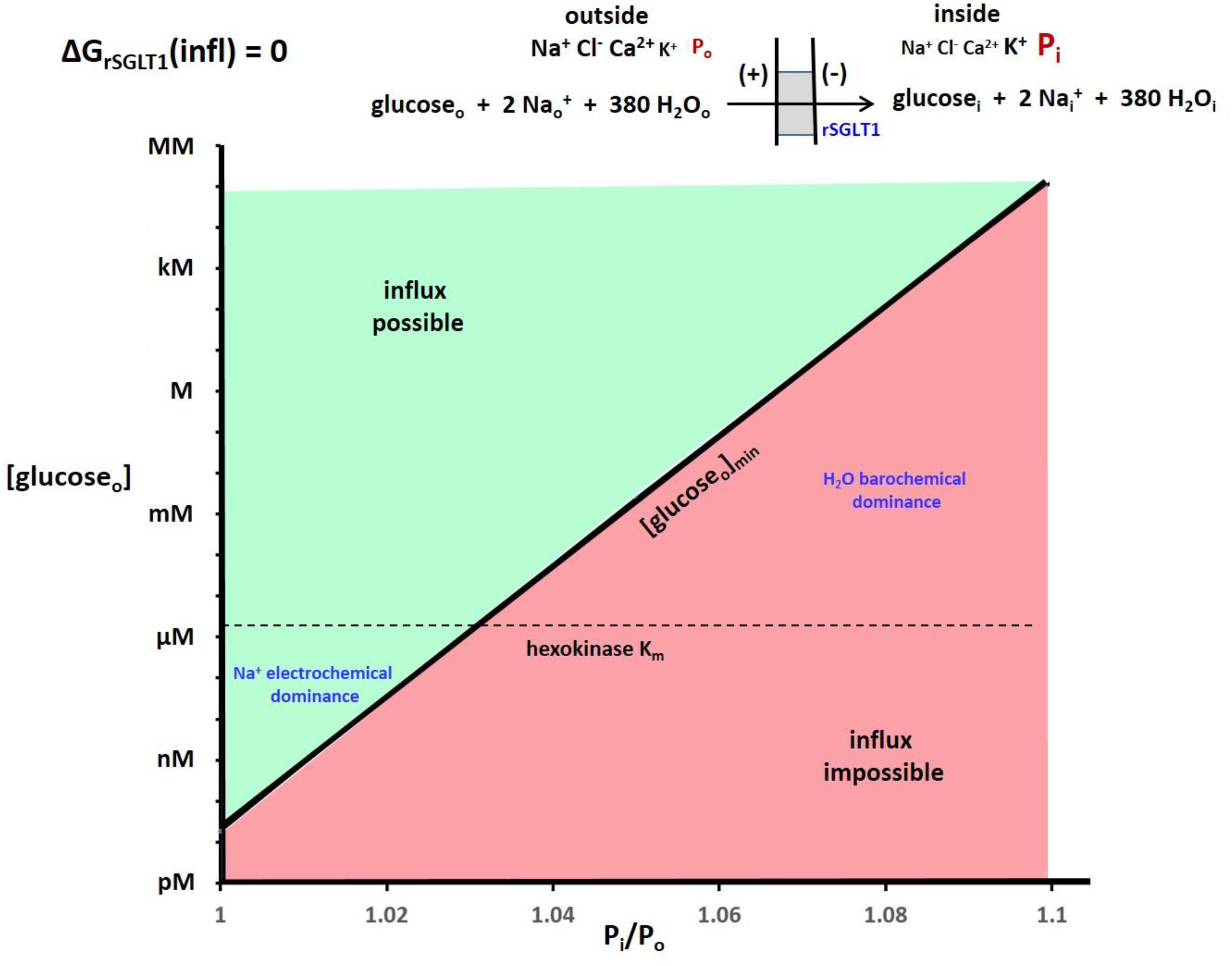
The 2D plot of the Figure 4 ΔG_rSGLT1_(influx) = 0 plane. The regions where the Na^+^ electrochemical gradient dominate and the H_2_O barochemical gradient dominate are indicated. The dependence is so strong that small percentage changes of P_i_ cause very large changes of the minimum [glucose_o_] required for glucose uptake. The hexokinase K_m_ for glucose_i_ (1.7 μM) is indicated with a horizontal line.

Plausible water chemogenic ΔG_H2O_(infl) valies (**-** 5 to 5 J/mole) are small compared with a typical barogenic magnitude (2.58ln(P_i_/P_o_) = 2.58ln(1.03) = 76 J/mole), which favors efflux. Thus, it is relatively inconsequential whether water is in chemical equilibrium (OsX_H2O,i_ = OsX_H2O,o_) or not. The larger steady-state water barochemical potential gradient is mostly barogenic.

Since it is likely almost all cells have P_i_ 1.01 atm or greater, it seems that each of the Table 3a reactions will have similar behaviors. The water barochemical contributions provide crucial counterbalances for other influx transporters as well.

Though the GLUT family of glucose influxers lacks the large SGLT Na^+^ electrochemical potential gradient favoring influx, it also has a smaller H_2_O barochemical force opposing influx when P_i_ > P_o_: the GLUT H_2_O stoichiometries are smaller (Fig. 1). **Figure 6** contrasts the 2D plot of GLUT1 with that of rSGLT1 (from Fig. 5) and makes obvious for the SGLT family the Na^+^ electrochemical dominance at small P_i_ values and the H_2_O barochemical dominance at large P_i_.

**Figure 6.**
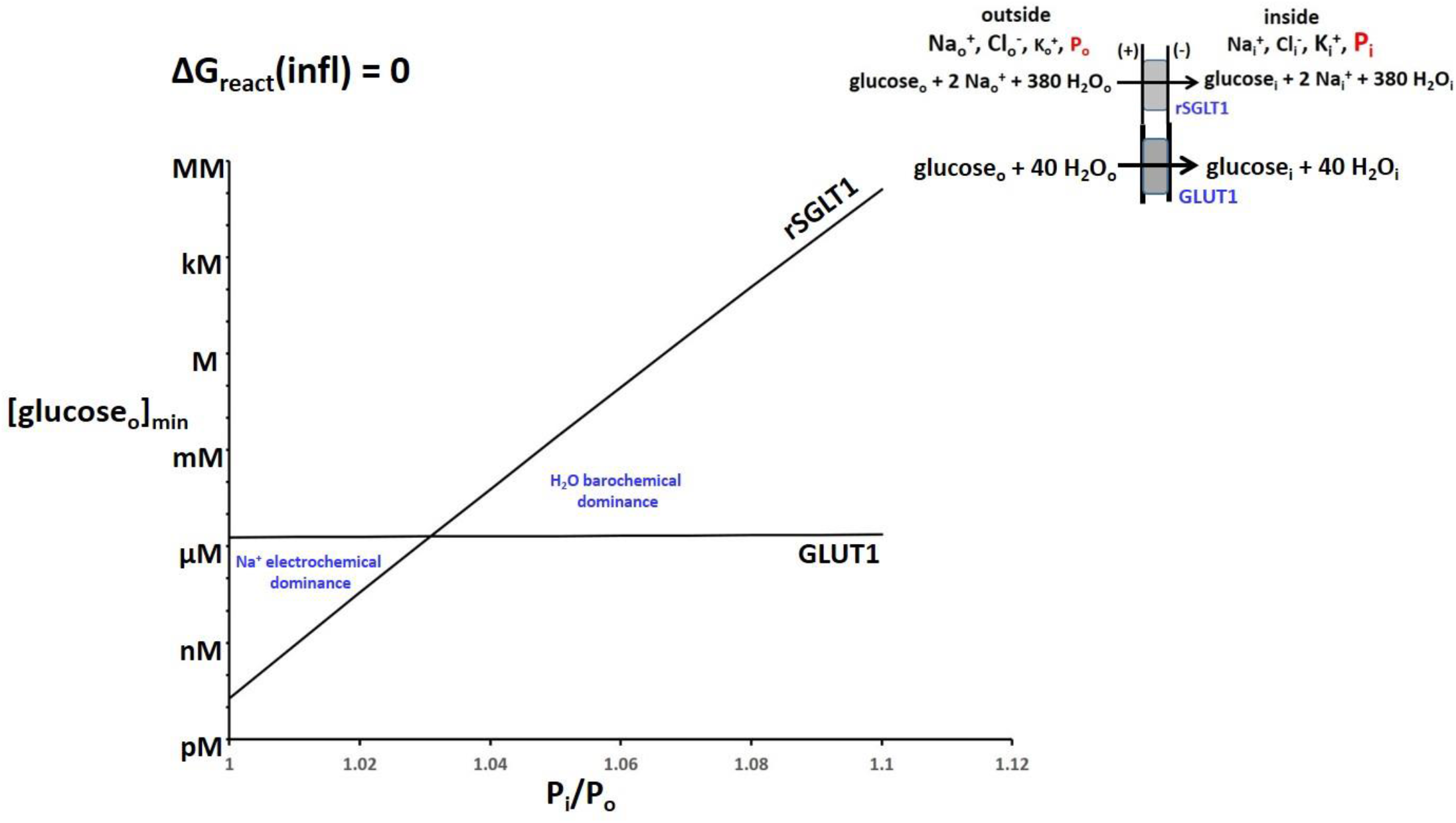
The (P_i_/P_o_)-dependences of [glucose_o_]_min_ for rSGLT1 (from Fig. 5) and for GLUT1. Equation (4) was used, with the intracellular glucose concentration, [glucose_i_], the membrane potential, E_m,oi_, and P_o_ held fixed at 2 μM, **-**91 mV, and 1 atm, respectively (T = 310 K). The regions of Na^+^ electrochemical and H_2_O barochemical dominance are very evident. Influx through the rSGLT1 transporter is much more sensitive to P_i_/P_o_ than that (barely noticeable) through the GLUT1 transporter. This is due to the much greater water stoichiometry of the former (Fig. 1).

##### Glucose Uptake at Fixed [glucose_o_]

For an alternative perspective, we take the extracellular glucose concentration, [glc_o_], to be maintained at 5 mM (a large, but reasonable (68,69), value). With rSGLT1, if we fix P_i_ = 1.05 P_o_ and neglect water co-transport, ΔG remains negative until the intracellular glucose concentration, [glc_i_], reaches the absurdly large value of 300 M. On the other hand, if we keep P_i_ = 1.05 P_o_ but retain the water co-transport, the reaction becomes unfavorable when [glc_i_] reaches only 4.1 μM. More generally for rSGLT1 with H_2_O transport, **Figure 7** shows the (P_i_/P_o_)-dependence of [glc_i_]_max_, with [glc_o_] fixed at 5 mM. The location of the hexokinase glucose_i_ K_m_ is shown as a horizontal dashed line. For such a plot, in the lower portion of the graph, the influx reaction is now possible (green), while it is impossible (red) in the upper portion. The maximum value of [glc_i_] is greatly suppressed by increasing P_i_/P_o_.

**Figure 7.**
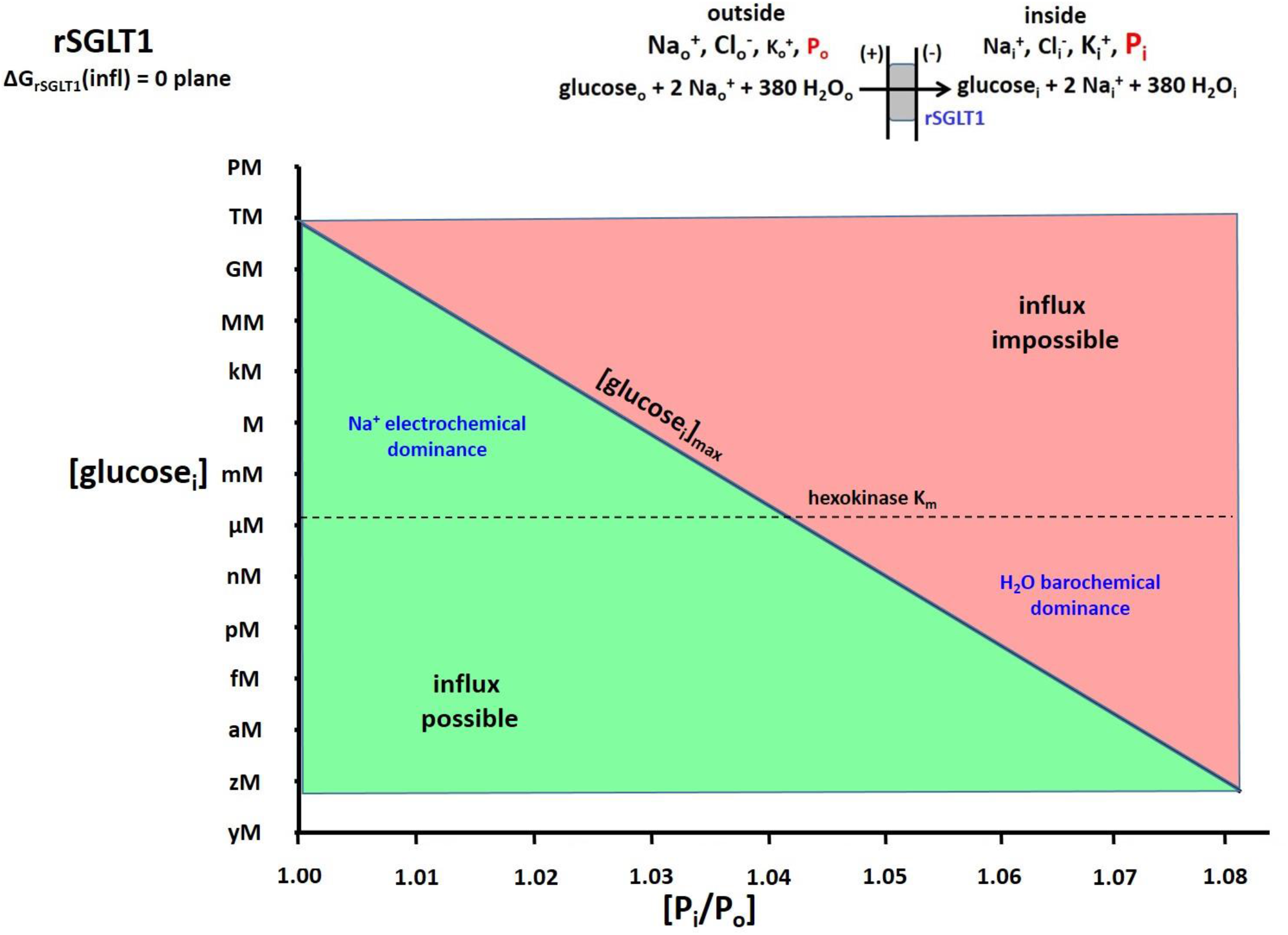
The 2D plot of the ΔG_rSGLT1_(influx) = 0 plane from a 3D plot similar to Figure 4. Equation (6) was used and, in this case, the *extracellular* glucose concentration, [glucose_o_], the membrane potential, E_m,oi_, and P_o_ were held fixed at 5 mM, **-**91 mV, and 1 atm, respectively (T = 310 K). The surface is colored green when influx is thermodynamically possible and red when it is impossible. Thus, the intersection of the ΔG surface with the ΔG = 0 plane traces the trajectory of the (P_i_/P_o_)-dependence of the (in this case) *maximum* intracellular glucose^**-**^ concentration, [glucose_i_]_max_, value allowing uptake. The K_m_ value of the cytoplasmic hexokinase for glucose_i_ (1.7 μM) is indicated with a horizontal dashed line. It is clear when P_i_ is small, tremendous values of [glucose_i_] are allowed, which would easily saturate hexokinase. However, this is not the case when P_i_ is only a few percentage points greater.

**Figure 8.**
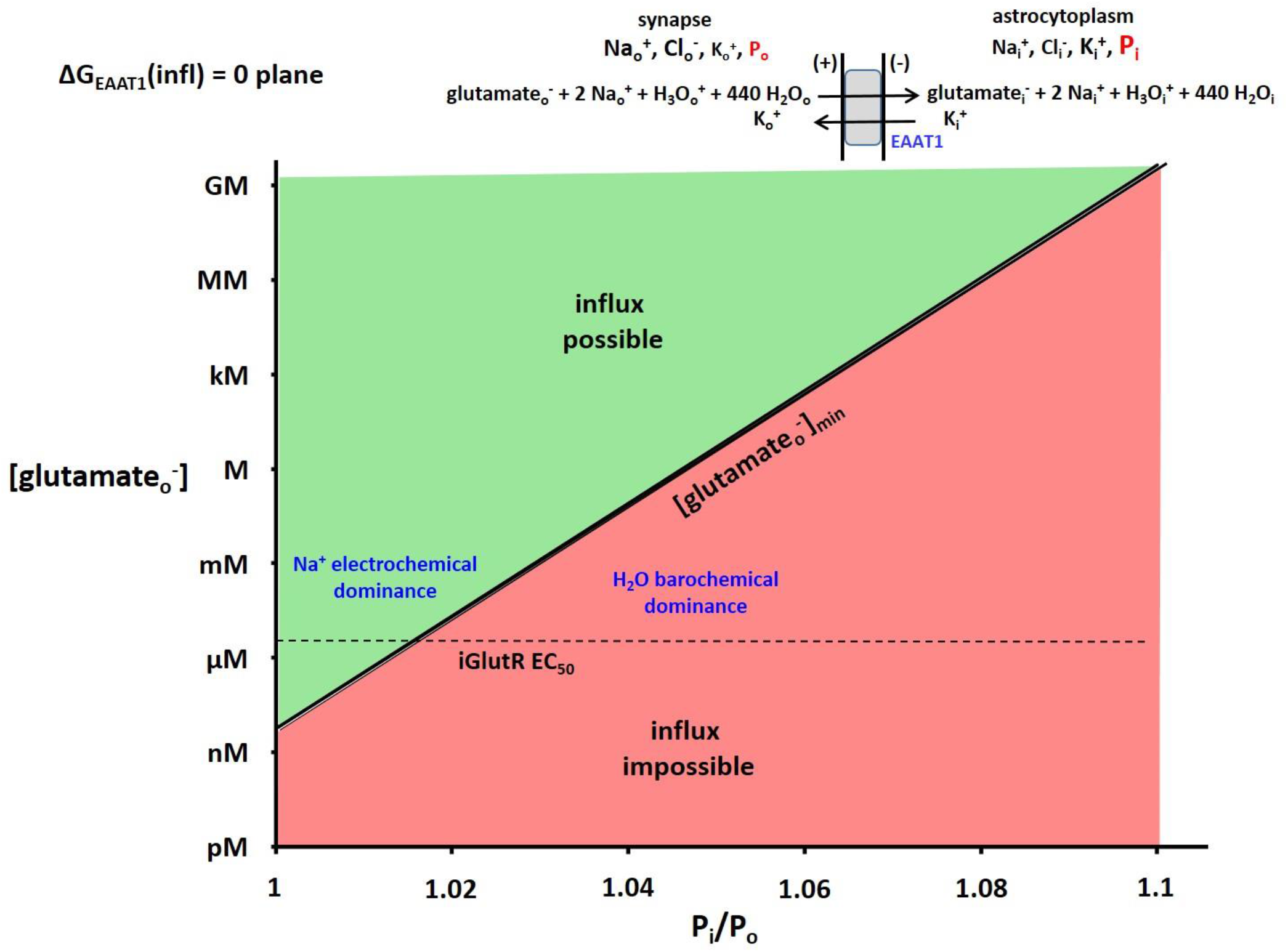
The 2D plot of the ΔG_EAAT1_(influx) = 0 plane from the 3D plot analogous to Figure 4. The (in this case) astrocytic glutamate^**-**^ concentration, [glutamate_i_^**-**^], the membrane potential, E_m,oi_, and P_o_ were fixed at 1.4 mM (the K_m_ value for glutamine synthetase), **-**91 mV, and 1 atm, respectively (T = 310 K). The surface is colored green when influx is thermodynamically possible and red when it is impossible. Thus, the intersection of the ΔG surface with the ΔG = 0 plane traces the trajectory of the (P_i_/P_o_)-dependence of the *minimum* synaptic glutamate^**-**^ concentration, [glutamate_o_^**-**^]_min_, value required for astrocyte uptake. The EC_50_ value of the iGlutR receptor for synaptic glutamate_o_^**-**^ (2.3 μM) is indicated with a horizontal dashed line. It is clear glutamate_o_^**-**^ is well-cleared from the synapse when P_i_ is small, and not when P_i_ is only a few percentage points greater. In the latter case, the receptors would be saturated, and synaptic transmission would be interrupted.

For GLUT2 at P_i_ = 1.04 P_o_ and [glc_o_] = 5 mM, [glc_i_] could accumulate to 4.8 mM if there was no water co-influx, but only 110 μM with H_2_O co-influx.

##### Neurotransmitter Uptake and Clearance

There are considerable similarities of the rSGLT and excitatory amino acid transporter (EAAT1) influx reactions – with the exceptions of the H_3_O^+^ co-influx and K_i_^+^ co-efflux of the latter. Glutamate^**-**^ influx is greatly aided by the Na^+^ electrochemical gradient, but opposed by the H_2_O barochemical gradient. The analogous EAAT1 3D plot (not shown) is very similar to that for rSGLT (Fig. 4). However, the roles of these two reactions are quite different. While the purpose of rSGLT is mainly to deliver glucose into cells, that of EAAT1 is principally to clear glutamate^**-**^ from synapses (*via* astrocytic uptake) after action potential transmission.

Now, besides E_m,oi_ = **-** 91 mV as before, we fix *intracellular* [glutamate_i**-**_] at 1.4 mM, the glutamate_i_^**-**^ K_m_ for glutamine synthetase (70), which converts astrocytic glutamate_i_^**-**^ to glutamine_i_. **Figure 8** depicts the 2D ΔG_EAAT1_(infl) = 0 plane of the unshown 3D plot. The vertical axis measures log *extracellular* [glutamate_o_^-^], while the horizontal axis plots P_i_/P_o_. The intersection of the ΔG_EAAT1_(infl) surface with the ΔG_EAAT1_(infl) = 0 plane traces the trajectory of the logarithm of the *minimum* glutamate_o_^**-**^ concentration ([glutamate_o_^**-**^]_min_) necessary to initiate influx, as a function of P_i_/P_o_. Though strictly bi-exponential, the plot is almost a single exponential function (the exponential nature of the P_i_/P_o_ axis is very weak). For a member of the ionotropic glutamate receptor (iGlutR) family, the position of the L-glutamate_o_^**-**^ EC_50_ value, 2.3 μM (71), is shown as a horizontal dashed line.

There is an influxer for which representative concentrations for all substrates are found in Table 2. K_o_^+^ influx *via* NKCC1 is not affected by the concentration of a metabolite substrate.

Inclusion of the effects of E_m,oi_ and ΔG_Na_ variation would increase the dimensionalities of Figure 4-type plots. Making ΔG_Na_(infl) less negative, for example, would (non-linearly) shift the lines to the left in Figs. 5-8. Making E_m,oi_ less negative (by increasing [K_o_^+^] or by [K_i_^+^] reduction) should do the same. In this paper, however, we focus on the novel (and large) effects of P_i_ variation, consequent to water co-transport.

##### Efflux Reactions

The ΔG_transporter_(effl) values for the normal *efflux* reactions (II, I, or both, in Figs. 1,2) are found in **Table 3b** (again, E_m,oi_ is fixed at – 91 mV). (Note: the calculations are for ΔG(effl), not ΔG(infl).) In contrast to the Table 3a influx reactions, except for that of KCC4 the other efflux reactions are unfavorable at P_i_/P_o_ = 1.00 (blue). But, they are made much more favorable as P_i_/P_o_ increases (red), because of water co-efflux.

There is an efflux reaction for which representative concentrations for all substrates are found in Table 2; KCC4, the [K^+^,Cl^**-**^,500H_2_O] effluxer. For this, we see the efflux favorability is greatly enhanced by water co-efflux, even though it is slightly favorable even w/o H_2_O co-transport. Thus, it will always operate in efflux mode. K_i_^+^ efflux *via* KCC4 is not limited by the concentration of a metabolite substrate.

We focus on the lactic acid exporter, MCT1. The reaction is unfavorable when P_i_/P_o_ = 1.00, ΔG_MCT1_(effl) = 8.5 kJ/mole (blue), but very favorable, **-** 58 kJ/mole (red), when P_i_/P_o_ = 1.05. Again, this is due to H_2_O co-efflux. At P_i_/P_o_ = 1.05 atm, the [lac_o_^**-**^]/[lac_i_^**-**^] ratio could reach only two without water, but can attain 3 x 10^10^ with the 500 H_2_O molecules co-effluxed.

## DISCUSSION

### Biochemical Roles for Water Co-Transport

The analyses above suggest co-transported water can exert significant thermodynamic effects on its process *via* the cytoplasmic pressure (P_i_). We elaborate some examples.

#### Figure 1 Glucose Influxers

##### Small [glucose_o_]

Commonly, the SGLT transporter family is thought of in terms of its potential for catalyzing glucose influx against its concentration gradient, chemogenic ΔG_glc_(infl) (72,73). However, Figs. 5 and 6 show this is true only when the P_i_ value is relatively small. For rSGLT1, “uphill” glucose influx is impossible for any P_i_ above ∼1.03 P_o_. A 1 % P_i_ increase to 1.04 atm requires a [glc_o_] value of at least ∼10 μM for influx. This is five times *larger than* the 2 μM [glc_i_] fixed for Figs. 5-6. Perhaps cells using exclusively SGLT transporters always have small P_i_ values. Maybe P_i_ fluctuation serves to control glucose influx. The GLUT family of transporters, with its smaller H_2_O stoichiometries (Fig. 1), does not suffer this severe (P_i_/P_o_)-dependence (Fig. 6). For GLUT1, the H_2_O/glucose transport ratio is much smaller than for rSGLT1, and its [glc_o_]_min_ value is almost P_i_-insensitive (Fig. 6). Perhaps cells with larger P_i_ values, and in situations with smaller [glc_o_] values, employ GLUT enzymes. On the other hand, if P_i_ values are small, cells with SGLT transporters will take up glucose at smaller [glc_i_] values than cells with GLUT transporters. When rSGLT1 P_i_ is small, small [glc_o_] can insure sufficient [glc_i_]. Also, when P_i_ is small, besides carrying glucose uphill, SGLT transporters are “pumping” water up a barochemical hill. They are maintaining the H_2_O barochemical steady-state.

##### Large [glucose_o_]

If there is a situation where extracellular glucose is maintained at a relatively large value (*e*.*g*., 5 mM), the role of P_i_ can be viewed differently. Setting P_i_ = P_o_, or ignoring co-transported water, would allow absurdly large [glc_i_]_max_ values for cells with SGLT enzymes (Fig. 7). This would surely amount to a sugar overload. Even if that was not cytotoxic, it could cause cells in a tissue to partake differentially of any available glucose charge – some cells initially reached by a bolus taking up more (or even all) sugar than others. This would mean a heterogeneous cellular [glc_i_] spatial distribution. However with H_2_O co-transport, representative experimental P_i_ values (say, 4% larger than P_o_) suppress the [glc_i_]_max_ value to near the hexokinase K_m_, 1.7 μM for glucose (Fig. 7). Cells with even larger P_i_ values will take up less glucose than those with smaller P_i_. For rSGLT1 when P_i_/P_o_ = 1.05 and [glc_o_] = 5 mM, [glc_i_]_max_ is near 1 nM, insufficient for metabolism. With no H_2_O co-transport, [glc_i_]_max_ would increase more than ten orders of magnitude; many orders greater than the hexokinase K_m_ for glucose.

For a given extracellular glucose concentration, cells with exclusively GLUT transporters will take up more glucose than cells with exclusively SGLT enzymes at larger P_i_ values. Perhaps GLUT transporters are found in cells with larger P_i_ values also in environments where [glc_o_] is large.

As shown in Fig. 1, both SGLT and GLUT transporters deliver glucose. It is interesting a switch from GLUT1-to SGLT-mediated cellular glucose uptake during lung cancer progression has been reported (20).

More generally, GLUT1 expression (along with other genetic changes) is found to promote the cancer cell Warburg State (74). However, among these families, only the GLUTs transport the ^18^FDG-PET tracer, (2-fluorodeoxyglucose) 2-FDG (20). This produces an interpretation problem for the metabolic rate of glucose uptake and consumption determined by quantitative ^18^FDG-PET.

Because of H_2_O co-transport, cells seem never allowed much steady-state free glucose. The role of water co-influx, common to all the glucose influxers [hSGLT1, rSGLT1, GLUT1, GLUT2], appears dominant in controlling [glc_i_]. It seems water co-transport *via* an SGLT generally guarantees a [glc_i_] near the hexokinase K_m_ for glucose. Glucose is effectively metabolized immediately upon entering the cell. This protects cells from too much glucose, and tissues from excessive cellular glucose uptake inequality. A large ([glc_o_] - [glc_i_]) difference makes [glc_i_] hard to measure, even with modern methods (47). Also, glc_i_ likely does not contribute to the glucoCEST NMR signal (75,76). On the other hand, GLUT1 generally keeps [glc_i_] near [glc_o_] (Fig. 6). (GLUT2 will keep it somewhat smaller.)

##### Figure 1 Neurotransmitter Influxers: astrocyte uptake / synaptic clearance

Similar considerations must also obtain for the Table 3a influx reactions of the principal excitatory and inhibitory neurotransmitters glutamate_o^**-**^_ and GABA_o_, respectively. Once inside the astrocyte, these must be processed immediately by their metabolizing enzymes. EAAT1 (Fig. 8) and GAT1 will not allow them to build-up. (Note: for EAAT1, Z_glt_ = - 1, and there are H_3_O_o_^+^ co-influx and K_i_^+^ co-*efflux* terms (21).)

However, an even more crucial aspect of the EAAT and GAT roles is the effective clearance of neurotransmitter species from synaptic junctions in time to enable the next action potential. Figure 8 shows this for EAAT1 and glutamate_o_^**-**^. It is clear the astrocyte P_i_ must be relatively small (< 1.02 atm) to ensure [L-glutamate_o_^**-**^] does not much exceed 2.3 μM, its EC_50_ value for the iGlutR receptor enzyme. At the same time, the astrocyte P_i_ must be large enough (1.035 atm) to ensure astrocytosolic [L-glutamate_i_^**-**^] does not much exceed 1.4 mM, its K_d_ value for glutamine synthetase (analogous to Fig. 7). The large H_2_O stoichiometries for EAAT1 (and GAT1) transport (similar to that of rSGLT1, Fig. 1) guarantee high sensitivity to astrocyte P_i_.

The astrocytic uptake of synaptic glutamate^**-**^ is particularly interesting. If astrocyte P_i_/P_o_ is even ∼1.03, [glutamate_o_^**-**^] would have to reach ∼100 μM (almost two orders of magnitude *over* 2.3 μM) to initiate uptake (Fig. 8). This could be sufficient to saturate iGlutR, and quench further synaptic transmission. Perhaps this is a physicochemical mechanism whereby astrocytes serve as “gatekeepers” for synaptic activity. Increases of P_i_ to only 3 or 4% above P_o_ could allow receptor saturation, and the interruption of neuronal firing. Very small astrocyte P_i_ fluctuations could enable or disable synaptic function. For excitatory glutamatergic synapses this means turning off or on neural *excitation*. A glance at the Fig. 1 GAT1 influx stoichiometry indicates very similar considerations would obtain for GABAergic synapses. In that case, neuronal *inhibition* could be halted by increased intra-astrocyte pressures – resulting in increased *excitation*.

Astrocyte swelling, a sign of possible P_i_ change, has been implicated in the glymphatic processes occurring during sleep (commentary (77)). Also, it has been suggested some therapeutic interventions can disturb the glutamate^**-**^/glutamine cycle leaving interstitial glutamine, which can act as a nutrient for brain cancer cells (78). In non-neural cells, [glutamate_i_^**-**^] values can become very large, 64 mM (47). Perhaps this is because there is no glutamine synthetase.

##### The Figure 1 NKCC1 Influxer

The function of NKCC1 (reviewed in (22)) is to effect cellular K_o_^+^ and H_2_O_o_ uptake against their respective chemical and barochemical gradients. Because of the huge H_2_O stoichiometry (Fig. 1), this process is predicted to be strongly P_i_-dependent.

##### Figure 1 Effluxers

For the Table 3b effluxers, each of the reactions are made significantly more favorable by increased intracellular pressure (only KCC4 is favorable when P_i_ = P_o_, and NaDC1 requires P_i_/P_o_ > 1.05). Again, this is due to the pressure effect on the free energy caused by the large numbers of co-transported H_2_O molecules.

We use the monocarboxylate transporter [MCT1] as an example. It’s main role is ridding the cell of lactic acid build-up from glycolytic-type metabolism. This is common in cancer cells (the Warburg Effect). Lactic acid can be sufficiently cytotoxic that MCT1 inhibition has been considered as a cancer therapy (79). **Equation (7)** expresses

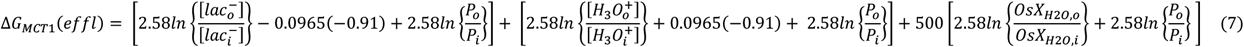

the free energy change for the lactic acid efflux reaction: the first, second, and third terms represent the lactate^**-**^, H_3_O^+^, and H_2_O contributions, respectively. Taking the Table 1 H_3_O^+^ and OsX_H2O_ concentrations, E_m,oi_ = -0.91 mV, {P_o_/P_i_} = {1/1.05} = 0.95, and letting ΔG_MCT1_(effl) = 0, we solve for the maximum {[lac_o_^-^]/[lac_i_^-^]} value that can be achieved. We find {[lac_o_^**-**^]/[lac_i_^**-**^]} = 2.6 x 10^10^ when 500 H_2_O molecules are co-transported, compared with only 2 if there was no H_2_O co-transport. This ten order-of-magnitude increase insures essentially all lactic acid is expelled from the cell – because of the barochemical water contribution. Once again, water co-transport is protecting the cells – even if they are malignant. A consequence is the extracellular acidification common in tumors (79). This phenomenon is also found for other cell types (*e*.*g*., muscle tissue cramping).

We now scrutinize the nature of intracellular hydraulic pressure (P_i_).

### The Nature of Intracellular Pressure

It seems clear the crucial thermodynamic role played by co-transported water is effectuated by the pressure difference across the cell membrane. The intracellular pressure (P_i_) is commonly considered “osmotic” in nature. That is, it is thought to result from the phenomenon of *water molecules* crossing the membrane. The simple first-order picture (H_2_O permeable; osmolytes (Na^+^, K^+^, Cl^**-**^) impermeable in the P_f_(p) sense) invokes *selective* water permeability, with influx and efflux becoming the same when ΔP (≡ P_i_ **-** P_o_) equals RT times the trans-membrane osmotic gradient (7): at T = 310 K, RT is 25.4 L•atm/mole. Strictly, osmotic effects are entropic in nature (the entropy of mixing (27)).

The first-order expression for the trans-cytolemmal osmotic gradient (Δπ_oi_) in terms of osmolyte concentrations is given in **Equation (8)**, where [osmolytes_i_] and [osmolytes_o_] are the intra- and extracellular concentrations,

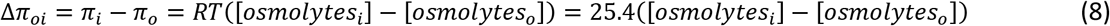

respectively. This equation assumes *ideal solutions* ((44), p 37; (80), p. 302 ff): *i*.*e*., entropy only. In Eq. (8), the osmolyte concentrations are given on the osmolarity scale. We use the representative values listed in Table 1. There, total [osmolytes_i_] = 0.379 OsM and total [osmolytes_o_] = 0.289 OsM. (At such magnitudes, these are essentially identical to their *molal* concentrations. We neglect the tiny mOsM intracellular RNA, lipid, and DNA concentrations.) Inserting these into Eq. (8) yields Δπ_oi_ = 2.29 atm. With the Table 1 compartmental water contents, we can also estimate volume molal (_V_m = OsM/f_W_) osmolyte concentrations. We obtain: _V_m_i_ = 0.38/0.76 = 0.50 mole(osmolytes)/L(cell water), and _V_m_o_ = 0.29/0.82 = 0.35 mole(osmolytes)/L(extracellular water). Inserting these into Eq. (8) gives: Δπ_oi_ = 25.4 (_V_m_i_ - _V_m_o_) = 25.4(0.50 – 0.35) = 3.8 atm, an even greater value.

If there were only first-order mixing entropy contributions, P_i_ would equal 2.3 or 3.8 atm at osmotic “equilibrium” (really steady-state; assuming P_o_ = 1 atm). A molecular mechanism for selective water influx and efflux could be free (*i*.*e*., unregulated) bidirectional aquaporin-mediated transport, as has been suggested (81). However, the largest experimental P_i_ value (1.5 atm (66)) is 1.5 times smaller than even 2.3 atm. The most pertinent P_i_ values (1.02 to 1.05; Figs. 4-8) are more than 2.2 times smaller. A (P_i_ - P_o_) of 1.03 would correspond to Δ[osmolytes] = 0.04 OsM (1.03/25.4). This is 2.5 times smaller than the 0.1 OsM value we estimate in Table 1. Of course, the Table 1 Δ[osmolytes] value is not found for any real cell. However, it is a reasonably representative magnitude: it is very unlikely to be less than 1.5 times any real value. If anything, 0.1 OsM may even be small (52). It is not clear any real cytoplasmic lipid bilayer membrane could even survive ΔP of 2.3 atm (the hydraulic pressure *ca*. 38 ft below sea level) if the *internal* pressure (P_i_) is the greater, let alone 3.8 atm. Even if it could, a ΔP value of even 2.29 atm would yield a barogenic Eq. (8) term: ΔG(infl) = 2.58ln(2.29) = 2.14 kJ/mole. This is so large, it would dominate the values in Tables 2 - 4, and obliterate the realistic estimations of the last section.

Such first-order considerations carry the implicit assumption of solution *ideality*; no specific osmolyte/osmolyte molecular interactions. Osmotic pressure is generally considered one of the “colligative” properties: the identity of the osmolyte is irrelevant. This is why the mole fraction concentration scale, mOsX (or the related mOsm scale), for solution components is the most appropriate for osmotic considerations ((44), p. 56; (45), p.27; (46)). But, since the Table 1 osmolyte mOSM values are sufficiently small (in the absolute sense) that mOsm values can be used, this suggests their activity coefficients (γ_osmolyte_, or y_osmolyte_) are near unity. However, such reasoning does “fail to emphasize the departure from ideality indicated by the activity of the solute” ((45), p.29).

Furthermore, we see from Table 1 intra- and extracellular water mole fractions OsX_H2O,i_ = 0.993 and OsX_H2O,o_ = 0.995, respectively. OsX_H2O,i_ is only 0.2% smaller than OsX_H2O,o_, and each is nearly the value for pure water. Given this situation, it is not unreasonable to also assume the water activity coefficients are equal, f_H2O,i_ = f_H2O,o_ (63). In **A.2**, we present calculations of H_2_O activities for surrogate interstitial and cytosolic solutions that support this contention. Table 2 shows inserting the calculated values a_H2O,i_ = 0.994 and a_H2O,o_ = 0.995 into Eq. (3) or (4), leads to a water chemogenic contribution to ΔG_H2O_(infl) of only ∼3 J/mole favoring cellular water influx. Also, we showed above that chemogenic ΔG_H2O_(infl) =**-** 0.003 kJ/mole is counteracted by a P_i_/P_o_ ratio of only 1.001. This is only one tenth of a conservatively small typical intracellular pressure, P_i_/P_o_ = 1.01 (27,66). This adds to the suggestion the entropic contribution to P_i_ is actually small, usually less than 10%.

All these considerations indicate the mixing entropic contribution (**-** TΔS_H2O_(infl)) to water ΔG_H2O_(infl) is small. By definition, a thermodynamically non-ideal solution means there is an enthalpic (ΔH_H2O_(infl)) contribution. A representative P_i_/P_o_ value of 1.05 corresponds to barochemical ΔG_H2O_(infl) = - 0.13 kJ/mole. A P_i_/P_o_ ratio of 1.001 implies ∼90% ΔH_H2O_(infl); *i*.*e*., - 0.12 kJ/mole. The conclusion must be that enthalpic (ΔH_H2O_(infl)) contributions dominate P_i_. The aqueous solutions inside (and, for that matter, outside) cells deviate greatly from ideality. There are highly specific molecular interactions between the various solutes, and with water. It has long been known the “hydration” of biological solutes is extremely important for many different cellular processes (57-59). If the solute-interacting water entropy is smaller than in pure water, this could even give rise to a (not mixing) entropic driving force for *efflux*. Whatever the actual case, the intracellular osmolytes do not seem to lower the water “escaping tendency” nearly as much as the first-order osmotic pressure equation (8) would predict.

There is a long history of considering higher order contributions to Eq. (8) ((82), p. 210 ff; (83)). However, it is probably not realistic to expect an environment as complex as the intracellular *milieu* to be well-modeled as a homogeneous solution.

Recent reports suggest particular intracellular species contribute to regulating intracellular hydraulic pressure. These include: the mechanosensitive transcriptional regulator YAP (Yes-associated protein) (84), tropomyosins 1.6 and 2.1 (Tpm 1.6 and Tpm 2.1) (85), and the capsaicin-activatable transient receptor potential vanilloid 1 (TRPV1) cation channel, that interestingly, stimulates the Fig. 1 NKCC1 water co-influxer (86) as well as NKA. An especially intriguing proposal is that intracellular macromolecular polyanion electrostatic interactions dominate P_i_, while allowing modulation by extracellular osmolality (52).

It may be often assumed tissue homeostasis means Δπ is zero, and is regulated to that point: *i*.*e*., [osmolytes_i_] = [osmolytes_o_] in Eq. (8): *i*.*e*., “*isosmolality*.” However, the word “*isotonal”* (6) suggests only equal pressures, and is preferred. A 290 mOsm NaCl solution is often considered “*isotonic*.” Bathing solutions hyper- or hypo-tonic relative to this cause *in vitro* cells (19) and *ex vivo* tissue cells (87) to shrink or swell due to *net* water efflux or influx, respectively. Bolus blood infusions of “*hypertonic*” solutions are used clinically to *transiently* open the blood-brain-barrier by shrinking capillary endothelial cells – and thus deliver otherwise non-extravasating therapeutic drugs to the cerebral parenchyma (88). In cell cultures and to some degree in perfused tissues, the investigator can specify and control [osmolytes_o_]. When such a system reaches homeostasis, a common presumption may be that [osmolytes_i_] = [osmolytes_o_]. From this, it is usually assumed [osmolytes_i_] ≈ 0.3 OsM, the Table 1 [osmolytes_o_] value (52). However, our estimated [osmolytes_i_] ≈ 0.4 OsM (Table 1) indicates this may be rarely (never) true.

Likely further evidence of cytoplasmic non-ideality is that different tissues seem to require different bathing solution osmolalities to achieve isotonicity. While ∼320 mOsm suffices for the retina and olfactory bulb, almost 600 mOsm is required for cerebral cortex (87). Inhibition of NKA causes cortical cells to swell considerably; *i*.*e*., a net category B water influx (89).

Whatever the actual P_i_ physicochemical nature, the calculations in the last section remain valid because they employ experimental P_i_ values.

### Aquaporin Role in Cellular Homeostasis

The specific water-transporting membrane aquaporin (AQP) molecules are found in almost all tissues (6,90,91). Like all the Fig. 1 transporters, they are capable of catalyzing category B unidirectional (flux, P_f_-type) or category A bidirectional (exchange, P_d_-type) water transport. The famously large “single channel” AQP4 water volume flux, 0.25 fL/s/AQP4 (8.3 H_2_O molecules/ns/AQP4), was determined at 10°C and with a very large 4.4 atm osmotic gradient (92). Like all P_f_ measurements, this is derived from the *asymptotic* volume change when transport is initiated away from osmotic “equilibrium.” However, AQP efficiency is minimal when there is no pressure gradient (ΔP = 0) (37), *i*.*e*., the P_d_-type condition. This is consistent with molecular considerations (the water single-file nature of the AQP channel structure (30) is not optimal for exchange). Thus, AQP’s by themselves have been thought of as intrinsically *passive* transporters. They can facilitate very large *net* water influxes or effluxes, but only when the latter are driven by independent forces. Thus, though their involvement in non-homeostatic cell swelling or shrinking has been widely investigated (6,90,91), their roles in homeostasis has been less clear. We are not aware a “single channel” category A exchange magnitude has ever been experimentally determined.

The results presented here suggest there is thermodynamic (metabolic) energy stored in tissue water compartmentalization itself, helping maintain the system in homeostasis. Thus, we have inserted AQPs explicitly into the active trans-membrane water cycling (AWC) scheme, rate-limited by ^c^MR_NKA_ (Fig. 2). Since AWC is, by definition, homeostatic, MR_H2O_(infl) = MR_H2O_(effl). However, it is unlikely AQP’s contribute equally to water influx and efflux. As stated above, if ΔP was actually osmotic in nature, unregulated dominant AQP activity would lead to ΔP values many times those measured. Thus, it is likely MR_AQP_(infl) ≠ MR_AQP_(effl).

In cell suspensions (and *in vivo*) astroglial aquaporin AQP4 expression has been shown to contribute to category A water exchange (81). Furthermore, studies with human C6 glioma (cancer) cell suspensions and TGN020 (a specific AQP4 inhibitor) are informative. The k_io_ rate constant ratios (^TGN020^k_io_)/(^C6^k_io_) is 0.69, while (^ouabain^k_io_)/(^C6^k_io_) is 0.65 (81). Inhibiting water exchange with extracellular TGN020 reduces k_io_ to the same extent as inhibiting it with extracellular ouabain, the specific NKA inhibitor. This indicates AQP4 contributes to AWC. Cancer cells in suspension may not be in the Warburg state (2,4).

The metabolite influxers (Fig. 1) can provide significant MR_H2O_(infl) values. The cellular glucose consumption rate, MR_glc_(consump), has been determined to be 38 μM(glc)/s/cell for murine kidney epithelial cells (47) and for brain tissue cells, assuming reasonable cell density and volume values (68). The glucose influx rate, MR_glc_(infl) must be at least MR_glc_(consump). But GLUT1 and rSGLT1 have different stoichiometries (Fig. 1). Thus, MR_H2O_(infl) would range from 1.8×10^9^ to 17×10^9^ H_2_O/s/cell for 2pL cells with, respectively, exclusively GLUT1 or exclusively rSGLT1 transporters. Using the ratio of the glutamate^**-**^-to-glutamine conversion flux, MR_glu-gln_(cycle) to MR_glc_(consump) (93), we estimate MR_H2O_(infl) provided by the EAAT1 transporter to be 4 x 10^7^ H_2_O/s/cell for a 2 pL astrocyte. This would be in addition to MR_H2O_(infl) provided by a glucose influxer.

Some so-called “metabolic water” is generated within the cell after glucose is imported. However, there are at most six H_2_O molecules produced per glucose molecule metabolized, depending on the glycolysis and oxidative phosphorylation proportions (94). This is small when compared with even the smallest Fig. 1 glucose influxer H_2_O stoichiometries.

Our most recent estimate of brain x, the MR_H2O_(AWC)/MR_NKA_ ratio, is > 10^6^ (2). We opined “such large stoichiometries suggest aquaporin participation in active trans-membrane water cycling.” Given the large cellular water influx rates, and the P_i_ > P_o_ pressure gradient, it seems likely MR_AQP_(effl) > MR_AQP_(infl). That is, in homeostasis AQP is likely mainly catalyzing water *efflux*. It is working in parallel with the KCC effluxer. This is why we indicate the inhibitor blocking MR_AQP_(effl) in Fig. 2.

Water should be considered a substrate for active trans-membrane water cycling. It has thermodynamic consequences. If it turns out NKA itself has its own water stoichiometry, AQPs could be then considered as also secondary active enzymes. If not, they are only tertiary active transporters – sharing water as a substrate with only the secondary active water co-transporters.

An interesting analogy with potassium ion-specific channels (PCs) (reviewed in (29); (95)) might obtain. It is thought the formally “passive” K_i_^+^ efflux through PCs dominates the production of E_m,oi_. The idea is K_i_^+^ efflux driven by the chemogenic term proceeds until the latter is balanced by the electrogenic term (Table 2), which actually requires very few effluxed K^+^ ions. After that, an “electrochemical” steady-state condition is reached, and *homeostatic* K_i_^+^ efflux is synchronized with ^c^MR_NKA_. If it is water transport that dominates the P_i_ value, perhaps there is an analogous H_2_O_i_ efflux driven by the barochemical gradient. This would tend to decrease P_i_. However, diminished P_i_ would allow the Na^+^ electrochemical gradient a greater role. This would tend to increase the water co-influxes of the many transporters that also bring in Na^+^ (*e*.*g*., NKCC- and SGLT-catalyzed influxes; Figs. 1-8), which would tend to increase P_i_. This “feedback loop” would then establish an NKA-maintained *steady-state*, and AQP-catalyzed water efflux is also synchronized with ^c^MR_NKA_. Thus, we have AQP playing a vital role in active trans-membrane water cycling (Fig. 2).

A trans-membrane pressure difference could also affect transporter *kinetics*. This is discussed in **A.3**.

## CONCLUSION

The Na^+^,K^+^-ATPase (NKA) enzyme function has long been known to maintain Na^+^ and K^+^ in trans-membrane electrochemical *steady-states*, which are far from chemical *equilibria* ([Na_i_^+^] = [Na_o_^+^], [K_i_^+^] = [K_o_^+^]). Here, we find metabolic energy released by NKA-catalyzed ATP hydrolysis is also required to maintain (and is thus stored in) in a trans-cytolemmal water barochemical steady-state. This is an inherent part of NKA function. An important difference from the Na^+^ and K^+^ cases is the trans-membrane water distribution is in (or near) chemical equilibrium ([H_2_O_i_] ≈ [H_2_O_o_]). Crucially, the barochemical H_2_O steady-state is significant, and enzymatic water co-transport has very important thermodynamic metabolic consequences. Active trans-membrane water cycling does not represent a futile cycle.

## ACKNOWLEDGEMENTS

The authors especially thank Professor Daniel Zuckerman for critical readings of manuscript drafts and very insightful suggestions, and Dr. August George for accomplishing initial calculations. They also thank Professors Joseph Ackerman, Valerie Anderson, James Balschi, James Bassingthwaighte, Cynthia Burrows, Deborah Burstein, Kieran Clarke, Ira Cohen, Michael Grabe, Fahmeed Hyder, Joanne Ingwall, Christopher Kroenke, Kenneth Krohn, Philip Kuchel, Xin Li, Daniel Liefwalker (for ref. (74)), Jeffrey Maki, Silvia Mangia, Richard Mathias, Anusha Mishra, William Rooney, Douglas Rothman, Suzanne Scarlatta, William Skach, and Mark Woods, and Drs. Joseph Armstrong, Mauro DiNuzzo, Bassam Haddad, Audrey O’Connor, and Gregory Wilson, and Messers. Eric Baetscher, Eric Baker, Brendan Moloney, and Joshua Schlegel for stimulating discussions.

## AUTHOR ROLES

**CSS** conceived the approach, developed the theory, carried out the calculations, and drafted the manuscript.

**MMP** edited the manuscript drafts, with particular attention to the metabolic aspects.

**TMB** contributed calculations and edited the manuscript drafts, especially scrutinizing the theory.

## Appendix A.1. Acronyms and Symbols

⟨A⟩ tissue or voxel mean cell surface area

AQP aquaporin

AQP4 aquaporin 4

AQP4KO aquaporin 4 knock-out

ATP adenosine triphosphate

ATP_i_ intracellular ATP

AWC active trans-membrane water cycling

a_e_ ρ⟨A⟩

a_H2O_ water thermodynamic activity

CEST chemical exchange saturation transfer

ΔG Gibbs free energy change

ΔG^0^ Standard ΔG

ΔG_YYY_ ΔG for substrate, enzyme, reaction; YYY

ΔG_YYY_(effl) ΔG for YYY efflux

ΔG_YYY_(infl) ΔG for YYY influx

ΔH_YYY_(infl) enthalpy change for YYY influx

Δπ_oi_ trans-cytolemmal osmotic gradient (in – out)

ΔS_YYY_(infl) entropy change for YYY influx

ΔV^**††**^ activation volume

d density (mass/volume)

EAAT1 excitatory amino acid transporter 1

E_m,oi_ trans-membrane electrical potential (in – out)

effl an efflux process

FDG 2-deoxy-2-^18^Fluoro-D-glucose

f_H2O_ water activity coefficient on the mole fraction scale (X)

f_W_ tissue water volume fraction

f_W,i_ intracellular f_W_ f_W,o_ extracellular f_W_

ϕ osmotic coefficient

G Gibbs free energy

GABA γ-amino butyric acid

GAT1 GABA transporter 1

GLUT1 glucose transporter 1

glc glucose

gln glutamine

glu glutamate^**-**^

γ_H2O_ water activity coefficient on the molality scale (m)

^1^H_2_O water proton MR signal

H_2_O_i_ intracellular water molecule

H_2_O_o_ extracellular water molecule

H_3_O_i_^+^ intracellular hydronium ion

H_3_O_o_^+^ extracellular hydronium ion

iGlutR an ionotropic glutamate^**-**^receptor

infl an influx process

K_i_^+^ intracellular K^+^

K_o_^+^ extracellular K^+^

KCC4 potassium, chloride transporter 4

]k kinetic rate constant

k_io_ cellular water efflux k (1/τ_i_)

k_io_(a) active k_io_ contribution

k_io_(p) passive k_io_ contribution

k_oi_(p) passive cellular water influx k contribution

M molarity concentration scale

MADI metabolic activity diffusion imaging

MCT1 mono-carboxylate transporter

MD molecular dynamics

MRI magnetic resonance imaging

MR_YYY_ YYY metabolic rate

MR_glu_(consump) MR of glucose consumption

MR_glu_(infl) MR of glucose influx

MR_Glu-gln_(cycle) MR of glutamate^**-**^-glutamine cycling

MR_O2_ MR of O2 consumption

^c^MR_YYY_ cellular metabolic rate for YYY process = ^t^MR_YYY_/ρ

^t^MR_YYY_ tissue metabolic rate for YYY process = ρ^c^MR_YYY_

MW_H2O_ molecular mass (“weight”) of water

m molality concentration scale

mOsM milli-osmolarity concentration scale

mOsm milli-osmolality concentration scale

mOsX milli-osmole fraction concentration scale

μ_YYY_ chemical potential of YYY; (∂G/∂n_YYY_)_T,P,n(≠ nYYY)_

Na_i_^+^ intracellular Na^+^

Na_o_^+^ extracellular Na^+^

NKA Na^+^,K^+^-ATPase (sodium pump)

NKCC1 sodium, potassium, chloride transporter 1

NMR nuclear magnetic resonance

n number of moles

OsM osmolarity concentration scale

OsM_YYY_ YYY OsM

Osm osmolality concentration scale

Osm_YYY_ YYY Osm

OsX osmole fraction concentration scale

OsX_YYY,i_ intracellular YYY OsX

OsX_YYY,o_ extracellular YYY OsX

P hydraulic (mechanical) pressure

P^0^ standard state P

P_d_ diffusional permeability coefficient (exchange)

P_d_(p) passive P_d_

P_dYYY_ YYY P_d_

P_f_ flux (flow) permeability coefficient

P_fYYY_ YYY Pf

P_i_ intracellular P

P_o_ extracellular P

PC potassium channel

PET positron emission tomography

p tissue water mole fraction (“population”)

p_i_ intracellular p

p_o_ extracellular p

per permeant particle

R ideal gas constant

**ρ** cell (number) density

S_YYY_ stoichiometric coefficient of YYY

SGLT sodium/glucose co-transporter

SS trans-cytolemmal NMR shutter-speed

TGN020 specific AQP4 inhibitor

τ_i_ mean H_2_O_i_ molecule lifetime (1/k_io_)

⟨V⟩ tissue or voxel mean cell volume

_V_m volume molality concentration scale

_V_m_i_ intracellular _V_m

_V_m_o_ extracellular _V_m

v tissue volume fraction

v_e_ extracellular v (ECV) (= 1 – v_i_)

v_i_ intracellular v [**ρ**V]

X mole fraction concentration scale

X_YYY_ YYY X

X_YYY,i_ intracellular YYY X

X_YYY,o_ extracellular YYY X

x water cycling stoichiometry [H_2_O/ATP]

x′ x for individual water co-transporter

[YYY] YYY concentration

[YYY_c_] compartmental YYY concentration

[YYY_i_] intracellular YYY concentration

[YYY_o_] extracellular YYY concentration

[YYY_t_] tissue YYY concentration = v_i_[YYY_i_] = v_e_[YYY_o_]

yL yocto liter (= 10^−24^ L = 1 (nm)^3^))

y_H2O_ water activity coefficient on the molarity scale (M)

Z_YYY_ signed electrical charge of YYY

### A.2. Compartmental Water Thermodynamic Activities

Solvent mole fraction values as large as OsX_H2O,o_ = 0.995 and OsX_H2O,i_ = 0.993 (Table 1) “fail to emphasize the departure from ideality indicated by the activity coefficient of the *solute*” ((45), p.29). However, here the *solvent* activity coefficient difference is also very small. Since the water activity a_H2O_ (dimensionless on this scale) is always so close to unity, it requires sophisticated apparatus for high-precision direct water vapor pressure determinations to measure (45). Using these, it has been found a_H2O_ is inversely, and exponentially, related to the solute osmolality, Osm_solute_, *via* the empirical *osmotic coefficient*, φ, **Equation (A.2.1)** (19,45,46; (63), p.12). MW_H2O_ is the water molecular

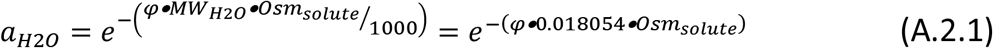

weight (mass), 18.0154 g/mole(H_2_O). The dimensionless product (MW_H2O_•Osm_solute_/1000) is the solute/solvent mole ratio (1000 is (g/kg)), and essentially the solute mole fraction, OsX_solute_. The latter is also dimensionless, and since the exponent must be dimensionless, φ is also dimensionless. Equation (A.2.1) provides a way to evaluate a_H2O_. (When Osm_solute_ is zero (pure water), a_H2O_ is exactly one.)

How might we more fruitfully evaluate Eq. (4) when per = H_2_O? Solutions as complex as those in Table 1 have never been subjected to precise measurements such as those of the last paragraph. However, pure NaCl and KCl solutions have been so studied, at 298 K ((45), p.476). We could take as a surrogate interstitium 0.290 Osm NaCl, and a surrogate cytosol 0.375 Osm KCl. (A glance at the metabolomics study mentioned above (47) indicates a K^+^glutamate^**-**^ solution would be a better intracellular surrogate. However, that has never been studied.) Their respective osmotic coefficients are *φ*_o_ = 0.9249 and *φ*_i_ = 0.9141 ((45), p.476) and, *via* Eq. (A.2.1), give a_H2O,o_ = 0.995 and a_H2O,i_ = 0.994. Using these activities in Eq. (4) gives ΔG(infl) = **-** 0.003 kJ/mole for H_2_O influx. This free energy change is again so small that correcting for temperature and actual solute content would make no difference. The conclusion is inescapable: there is a very small trans-membrane chemical potential difference (∼3 J/mole) for water to enter the cell.

### A.3. Intracellular Pressure and Kinetics (Reaction Activation Volumes)

Mechanical pressure can affect transporter free energies (as above), and macromolecular structures (96), but can also exert an effect on chemical reaction kinetics. This is characterized by the volume of activation, ΔV^**††**^, the difference between the partial molar volume of the reaction transition state and the partial molar volume of the reactants (97-99). The pressure dependence of a rate constant, k, is expressed in **Equations (A.3.1) and (A.3.2)**,

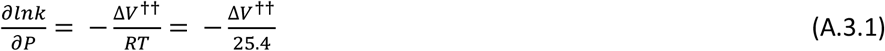

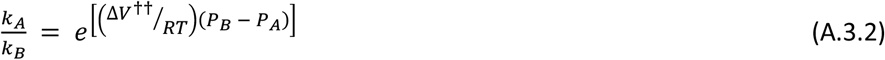

where RT is 25.4 L•atm/mole at 310 K. If a process has ΔV^**††**^ = 6 L/mole, a 10 atm pressure increase will cause its rate constant to decrease by 90%. Six L/mole is 10^**-**23^ L/molecule, or 10 yL/molecule (1 yocto liter = 1 yL = 10^**-**24^ L = 1 (nm)^3^). The plasma membrane NKA is a macromolecular complex of mass 145 kDa (145,000 g/mole). Thus (assuming d = 1 g/mL), it has a partial molar volume of approximately 242 yL/molecule. A 6 yL ΔV^**††**^ would represent only a 2.5 % volume change. For P_i_ = 1.5 atm (the maximum P_i_ reported), P_o_= 1 atm, and ΔV^**††**^ **=** 6 L/mole, Eq. (A.3.2) yields k_1.5 atm_ = 0.89 k_1 atm_, an 11% decrease.

It is easy to imagine ΔV^**††**^ values of this magnitude, or greater. In a molecular dynamics study of a small 6.5 kDa protein, Persson and Halle report a relatively constrained, 2 kDa domain undergoes numerous spontaneous 3% conformational volume fluctuations in a 1000 μs period (60). Most membrane transporters are rather large macromolecular complexes. It is quite likely they undergo significant absolute volume changes in reaching their transition states. Thus, their enzyme activities could be quite sensitive to intracellular pressure development. Some would slow down, and some would speed up: ΔV^**††**^ quantities can be negative (97-99). Furthermore, the activities of most membrane transporters are coupled to each other *via* common substrates or products (Fig. 1). Since the cell has evolved to have trans-membrane fluxes in homeostatic balance, it is likely a P_i_ change would disturb the latter. The consequent flux changes could serve an auto-regulatory function.

## REFERENCES

1. C. S. Springer, E. M. Baker, X. Li, B. Moloney, G. J. Wilson, M. M. Pike, T. M. Barbara, W. D. Rooney, J. H. Maki, Metabolic activity diffusion imaging [MADI]: I. Metabolic, cytometric modeling and simulations. NMR Biomed. 36, e4781 (2023).

2. C. S. Springer, E. M. Baker, X. Li, B. Moloney, G. J. Wilson, V. C. Anderson, M. K. Sammi, M. M. Pike, M. G. Garzotto, R. P. Kopp, F. V. Coakley, W. D. Rooney, J. H. Maki, Metabolic activity diffusion imaging [MADI]: II. Non-invasive, high-resolution human brain imaging of sodium pump flux and cell metrics. NMR Biomed. 36, e4782 (2023).

3. J. J. Neil, J. J. H. Ackerman, Metabolic activity diffusion imaging (MADI): A new paradigm. NMR in Biomed. 36, e4841 (2023).

4. J. Schlegel, E. Baker, S. Holland, J. Stoller, W. Packwood, X. Li, R. Barajas, C. Springer, M. Pike, Metabolic activity diffusion imaging [MADI] of rat brain glioma. Proc. Int. Soc. Magn. Reson. Med. 31, 3924 (2023).

5. M. M. Pike, X. Li, E. Baetscher, T. M. Barbara, M. K. Sammi, A. A. Stevens, C. S. Springer, Does MADI detect temporal brain metabolic activity changes?” Proc. Int. Soc. Magn. Reson. Med. 31, 5177 (2023).

6. R. E. Day, P. Kitchen, D. S Owen, C. Bland, L. Marshall, A. C. Conner, R. M. Bill, M. T. Conner, Human aquaporins: Regulators of transcellular water flow. Biochim. Biophys. Acta 1840, 1492–1506 (2014). [10.1016/j.bbagen201309033]

7. A. S. Verkman, Water permeability measurements in living cells and complex tissues. J. Membrane Biol. 173, 73–87 (2000).

8. E. Zeuthen, A sensitive ‘Cartesian Diver’ balance. Nature 159, 440–441 (1947).

9. O. Kedem, A. Katchalsky, Thermodynamic analysis of the permeability of biological membranes to non-electrolytes. Biochim. Biophys. Acta 27, 229–246 (1958).

10. C. R. House, Water transport in cells and tissues. (Edward Arnold, London, 1974), p. 156.

11. S-T. Chen, C. S. Springer, Ionophore-catalyzed cation transport between phospholipid inverted micelles manifest in DNMR. Biophys. Chem. 14, 375–388 (1981).

12. T. Conlon, R. Outhred, Water diffusion permeability of erythrocytes using an NMR technique. Biochim. Biophys. Acta 288, 354–361 (1972).

13. C. S. Landis, X. Li, F. W. Telang, P. E. Molina, I. Pályka, G. Vétek, C. S. Springer, Equilibrium transcytolemmal water-exchange kinetics in skeletal muscle in vivo. Magn. Reson. Med. 42, 467–478 (1999).

14. X. Li, W. D. Rooney, C. S. Springer, A unified magnetic resonance imaging pharmacokinetic theory: Intravascular and extracellular contrast reagents. Magn. Reson. Med. 54, 1351–1359 (2005). [10.1002/mrm.20684]

15. C. S. Springer, Using 1H2O to measure and map sodium pump activity in vivo. J. Magn. Reson. 291, 110–126 (2018). [10.1016/j/jmr.2018.02.018]

16. B. P. Hills, P. S. Belton, NMR studies of membrane transport. Ann. Reports NMR Spectroscopy 21, 99–159 (1989).

17. B. M. Denker, B. L. Smith, F. P. Kuhajda, P. Agre, Identification, purification, and partial characterization of a novel Mr 28,000 integral membrane protein from erythrocytes and renal tubules. J. Biol. Chem. 263, 15634–15642 (1988).

18. N. MacAulay, Molecular mechanisms of brain water transport. Nat. Rev. Neurosci. 22, 326–344 (2021).

19. T. Zeuthen, Water-transporting proteins. J. Membrane Biol. 234, 57–73 (2010). [10.1007/s00232-009-9216-y]

20. J. R. Barrio, S. C. Huang, N. Satyamurthy, C. S. Scafoglio, A. S. Yu, A. Alavi, K. A. Krohn, Does 2-FDG-PET accurately reflect quantitative in vivo glucose utilization. J. Nucl. Med. 61, 931–937 (2020).

21. N. MacAulay, U. Gether, D. A. Klaerke, T. Zeuthen, Water transport by the human Na+-coupled glutamate cotransporter expressed in Xenopus oocytes. J. Physiol. 530, 367–378 (2001).

22. K. Kaila, T. J. Price, J. A. Payne, M. Puskarjov, J. Voipio, Cation-chloride cotransporters in neuronal development, plasticity and disease. Nature Rev. Neurosci. 15, 637–654 (2014). [doi:10.1038/nrn3819]

23. A. W. Autry, J. W. Gordon, H-Y. Chen, M. LaFontaine, R. Bok, M. Van Criekinge, J. B. Slater, L. Carvajal, J. E. Villanueva-Meyer, S. M. Chang, J. L. Clarke, J. M. Lupo, D. Xu, P. E. Z. Larson, D. B. Vigneron, Y. Li, Characterization of serial hyperpolarized 13C metabolic imaging in patients with glioma. NeuroImage: Clin. 27, 102323 (2020).

24. R. Ye, A. S. Verkman, Simultaneous optical measurement of osmotic and diffusional water permeability in cells and liposomes. Biochem. 28, 824–829 (1989).

25. X. Li, S. Mangia, J-H. Lee, R. Bai, C. S. Springer, NMR shutter-speed elucidates apparent population inversion of 1H2O signals due to active transmembrane water cycling. Magn. Reson. Med. 82, 411–424 (2019). [doi: 10.1002/mrm.27725]

26. Y. Zhang, M. Poirier-Quinot, C. S. Springer, J. A. Balschi, Active trans-plasma membrane water cycling in yeast is revealed by NMR. Biophys. J. 101, 2833–2842 (2011). [doi:10.1016/j.bpj.2011.10.035]

27. Y. Li, K. Konstantopoulos, R. Zhao, Y. Mori, S. X. Sun, The importance of water and hydraulic pressure in cell dynamics. J. Cell. Sci. 133, jcs240341 (2020).

28. C. S. Springer, Measurement of metal cation compartmentalization in tissue by high resolution metal cation NMR. Ann. Rev. of Biophys. and Biophys. Chem., 16, 375–399 (1987).

29. C. Öster, K. Kendriks, W. Kopec, V. Chevelkov, C. Shi, D. Michl, S. Lange, H. Sun, B. L. de Groot, A. Lange, The conduction pathway of potassium channels is water free under physiological conditions. Sci. Adv. 5, eeaw6756 (2019).

30. B. L. de Groot, H. Grubmüller, Water permeation across biological membranes: Mechanism and dynamics of aquaporin-1 and GlpF. Science 294, 2353–2357 (2001).

31. R. Bai, C. S. Springer, D. Plenz, P. J. Basser, Brain active trans-membrane water cycling measured by MR is associated with neuronal activity. Magn. Reson. Med. 81, 1280–1295 (2019). [doi:10.1002/mrm.27473]

32. J. L. Adelman, C. Ghezzi, P. Bisignano, D. D. F. Loo, S. Choe, J. Abramson, J. M. Rosenberg, E. M. Wright, M. Grabe, Stochastic steps in secondary active sugar transport. Proc. Nat. Acad. Sci. 113, E3960–E3966 (2016).

33. J. L. Adelman, Y. Sheng, S. Choe, J. Abramson, E. M. Wright, J. M. Rosenberg, M. Grabe, Structural determinants of water permeation through the sodium-galactose transporter vSGLT. Biophys. J. 106, 1280–1289 (2014).

34. J. Li, S. A. Shaikh, G. Enkavi, P-C. Wen, Z. Huang, E. Tajkhorshid, Transient formation of water-conducting states in membrane transporters. Proc. Nat. Acad. Sci. 110, 7696–7701 (2013).

35. S. Choe, J. M. Rosenberg, J. Abramson, E. M. Wright, M. Grabe, Water permeation through the sodium-dependent galactose cotransporter vSGLT.” Biophys. J. 99, L56–L58 (2010).

36. A. Watanabe, S. Choe, V. Chaptal, J. M. Rosenberg, E. M. Wright, M. Grabe, J. Abramson, The mechanism of sodium and substrate release from the binding pocket of vSGLT. Nature 468, 988–991 (2010).

37. F. Zhu, E. Tajkhorshid, K. Schulten, Theory and simulation of water permeation in aquaporin-1. Biophys. J. 86, 50–57 (2004).

38. S. Zhang, J. Zhou, Y. Zhang, T. Liu, P. Friedel, W. Zhou, S. Somasekharan, K. Roy, L. Zhang, Y. Liu, X. Meng, H. Deng, W. Zeng, G. Li, B. Forbush, M. Yang, The structural basis of function and regulation of neuronal cotransporters NKCC1 and KCC2. Comm. Biol. 4, 226 (2021).

39. H. Ogawa, T. Shinoda, F. Cornelius, C. Toyoshima, Crystal structure of the sodium-potassium pump (Na+,K+-ATPase) with bound potassium and sodium.” Proc. Nat. Acad. Sci. 106, 13742–13747 (2009).

40. M. M. Pike, J. C. Frazer, D. F. Dedrick, J. S. Ingwall, P. D. Allen, C. S. Springer, T. W. Smith, 23Na and 39K nuclear magnetic resonance studies of perfused rat hearts. Biophys. J. 48, 159–173 (1985).

41. A. L. Lehninger, “Principles of Biochemistry.” (Worth Pub., New York, 1982), p. 571.

42. D. M. Bers, W. H. Barry, S. Despa, Intracellular Na+ regulation in cardiac myocytes. Cardiovas. Res. 57, 897–912 (2003).

43. A. A. Kadir, B. J. Stubbs, C-R, Chong, H. Lee, M. Cole, C. Carr, D. Hauton, J McCullagh, R. D. Evans, K. Clarke, On the interdependence of ketone body oxidation, glycogen content, glycolysis, and energy metabolism in the heart. J. Physiol. 601.7, 1207–1224 (2023). [doi:101113/JP284270]

44. A. G. Marshall, “Biophysical Chemistry: Principles, Techniques, and Applications” (Wiley, New York, 1978).

45. R. A. Robinson, R. H. Stokes, in “Electrolyte Solutions” (Dover, Mineola, ed. 2nd rev., 1970).

46. M. J. Blandamer, J. B. F. N. Engberts, P. T. Gleeson, J. C. R. Reis, Activity of water in aqueous systems: A frequently neglected property.” Chem. Soc. Rev. 34, 440–458 (2005).

47. J. O. Park, S. A. Rubin, Y-F. Xu, D. Amador-Noguez, J. Fan, T. Shlomi, J. D. Rabinowitz, Metabolite concentrations, fluxes, and free energies imply efficient enzyme usage. Nat. Chem. Biol. 12, 482–489 (2016).

48. O. S. Andersen, “Cellular electrolyte metabolism,” in Enclycopedia of Metalloproteins, R. H. Kretsinger, V. N. Uversky, A. Permiakov, Eds. (Springer, 2013), pp. 580–587.

49. R. Milo, What is the total number of protein molecules per cell volume? A call to rethink some published values. Bioessays. 35, 1050–1055 (2013). [10.1002/bies.201300066]

50. F. C. Neidhardt, H. E. Umbarger, in Escherichia coli and Salmonella: Cellular and Molecular Biology, F. C. Neidhardt, Ed. (Am. Soc. Microbiol. Press, New York, ed. 2, 1996), vol. 1, chapt. 3.

51. K. C. Vinnakota, J. B. Bassingthwaighte, Myocardial density and composition: A basis for calculating intracellular metabolite concentrations. Am. J. Physiol. Heart Circ. Physiol. 286, H1742–H1749 (2004). [10.1152/ajpheart.00478-2003]

52. H. Wennerström, M. Oliveberg, On the osmotic pressure of cells. QRB Discovery. 3, e12 (2022).

53. R. Milo, R. Phillips, N. Orme, “Cell Biology By the Numbers.” (Garland Science, New York, 2016), pp. 70, 92, 93, 96, 106, 130, 198.

54. M. A. Model, E. Schonbrun, Optical determination of intracellular water in apoptotic cells. J. Physiol. 591.23, 5843–5849 (2013). [doi: 10.1113/jphysiol.2013.263228]

55. W. D. Rooney, G. Johnson, X. Li, E. R. Cohen, S-G. Kim, K. Ugurbil, C. S. Springer, Magnetic field and tissue dependencies of human brain longitudinal 1H2O relaxation in vivo. Magn. Res. Med. 57, 308–318 (2007).

56. R. P. Rand, Probing the role of water in protein conformation and function. Phil. Trans. R. Soc. Lond. B 359, 1277–1285 (2004). [10.1098/rstb.2004.1504]

57. P. Ball, Water is an active matrix of life for cell and molecular biology. Proc. Nat. Acad. Sci. 114: 13327-13335 (2017). [10.1073/pnas.1703781114]

58. M. Chaplin, Do we underestimate the importance of water in biology. Nat. Rev. Mol. Cell Biol. 7, 862–866 (2006).

59. K. D. Garlid, The state of water in biological systems. Int. Rev. Cytology 192, 281–302 (2000).

60. F. Persson, B. Halle, Transient access to the protein interior: Simulation versus NMR. J. Am. Chem. Soc. 135, 8735–8748 (2013). [10.1021/ja403405d]

61. X. L. Lu, V. C. Mow, Biomechanics of articular cartilage and determination of material properties. Med. Sci. Sports Exerc. 40, 193–199 (2008).

62. E. Nimer, R. Schneiderman, A. Maroudas, Diffusion and partition of solutes in cartilage under static load. Biophys. Chem. 106, 125–146 (2003). [doi:10.1016/S0301-4622(03)00157-1]

63. K. S. Pitzer, in “Activity Coefficients in Electrolyte Solutions” (CRC Press, Boca Raton, ed. 2, 1991).

64. J. Gao, X. Sun, L. C. Moore, T. W. White, P. R. Brink, R. T. Mathias, Lens intracellular pressure is generated by the circulation of sodium and modulated by gap junction coupling. J. Gen. Physiol. 137, 507 520 (2011).

65. R. J. Petrie, H. Koo, Direct measurement of intracellular pressure. Curr. Protoc. Cell Biol. 63, 12.9.1-12.9.9 (2015).

66. J. Gao, X. Sun, T. W. White, N. A, Delamere, R. T. Mathias, Feedback regulation of intracellular hydrostatic pressure in surface cells of the lens. Biophys. J. 109, 1830–1839 (2015).

67. E. LoCastro, R. Paudyal, Y. Mazaheri, V. Hatzoglou, J. H. Oh, Y. Lu, A. S. Konar, K. vom Eigen, A. Ho, J. R. Ewing, N. Lee, J. O. Deasy, A. Shukla-Dave, Computational modeling of interstitial fluid pressure and velocity in head and neck cancer based on dynamic contrast-enhanced magnetic resonance imaging: Feasibility analysis. Tomog. 6, 129–138 (2020).

68. M. DiNuzzo, G. A. Dienel, K. L. Behar, O. A. Petroff, H. Benveniste, F. Hyder, F. Giove, S. Michaeli, S. Mangia, S. Herculano-Houzel, D. L. Rothman, Neurovascular coupling is optimized to compensate for the increase in proton production from nonoxidative glycolysis and glycogenolysis during brain activation and maintain homeostasis of pH, pCO2, and pO2. J. Neurochem. 00, 1–31 (2023). [doi:10.1111/jnc.15839]

69. H. H. Chowdhury, Differences in cytosolic glucose dynamics in astrocytes and adipocytes measured by FRET-based nanosensors. Biophys. Chem. 261, 106377 (2020).

70. I. Schomburg, A. Chang, S. Placzek, C. Sohnigen, M. Rother, M. Lang, C. Munaretto, S. Ulas, M. Stelzer, A. Grote, M. Scheer, D. Schomberg, BRENDA in 2013: Integrated reactions, kinetic data, enzyme function data, improved disease classification: New options and content in BRENDA. Nucleic Acids Res. 41, D764–D772 (2013).

71. D. K. Patneau, M. L. Mayer, Structure-activity relationships for amino acid transmitter candidates acting at N-Methyl-D-aspartate and quisqualate Receptors. J. Neurosci. 10, 2385–2399 (1990).

72. P. Bisignano, C. Ghezzi, H. Jo, N. F. Polizzi, T. Athoff, C. Kalyanaraman, R. Friemann, M. P. Jacobson, E. M. Wright, M. Grabe, Inhibitor binding mode and allosteric regulation of Na+-glucose symporters. Nature Comm. 9, 5245 (2018).

73. J. Jurcovicova, Glucose transport in brain – Effect of inflammation. Endo. Reg. 48, 35–48 (2014). [doi:10.4149/endo_2014_01_35]

74. J-H. Lee, R. Liu, J. Li, Y. Wang, L. Tan, X-J. Li, X. Qian, C. Zhang, Y. Xia, D. Xu, W. Guo, Z. Ding, L. Du, Y. Zheng, Q. Chen, P. L. Lorenzi, G. B. Mills, T. Jiang, Z. Lu, EGFR-phosphorylated platelet isoform of phosphofructokinase 1 promotes PI3K activation. Mol. Cell 70, 197–210 (2018).

75. X. Xu, A. A. Sehgal, N. N. Yadav, J. Laterra, L. Blair, J. Blakeley, A. Seidemo, J. M. Coughlin, M. G. Pomper, L. Knutsson, P. C. M. van Zijl, D-glucose weighted chemical exchange saturation transfer (glucoCEST)-based dynamic glucose enhanced (DGE) MRI at 3T: Early experience in healthy volunteers and brain tumor patients. Magn. Reson. Med. 84, 247–262 (2020).

76. F. A. Nasrallah, G. Pagès, P. W. Kuchel, X. Golay, K-H. Chaung, Imaging brain deoxyglucose uptake and metabolism by GlucoCEST MRI. J. Cerebr. Blood Flow & Metabol. 33, 1270–1278 (2013).

77. A. J. Smith, B-J. Jin, A. S. Verkman, Muddying the water in brain edema? Trends Neurosci. 38, 331–332 (2015).

78. T. N. Seyfried, L. Shelton, G. Arismendi-Morillo, M. Kalamian, A. Elsakka, J. Maroon, P. Mukherjee, Provocative question: Should ketogenic metabolic therapy become the standard of care for glioblastoma? Neurochem. Res. 44, 2392–2404 (2019).

79. K. Nath, D. S. Nelson, D. F. Heitjan, R. Zhou. D. B. Leeper, J. D. Glickson, Effects of hyperglycemia on lonidamine-induced acidification and de-energization of human melanoma xenografts and sensitization to melphalan. (2015). NMR Biomed. 28, 395-403. [DOI: 10.1002/nbm.3260]

80. H. B. Callen, “Thermodynamics and an introduction to thermostatistics.” (Wiley, New York, ed. 2, 1985).

81. Y. Jia, S. Xu, G. Han, B. Wang, Z. Wang, C. Lan, P. Zhao, M. Gao, Y. Zhang, W. Jiang, B. Qiu, R. Liu, Y-C. Hsu, Y. Sun, C. Liu, Y. Liu, R. Bai, Transmembrane water-efflux rate measured by magnetic resonance imaging as a biomarker of the expression of aquaporin-4 in gliomas. Nat. Biomed. Eng. 7, 236–252 (2023).[10.1038/s41551-022-00960-9]

82. C. Tanford, “Physical Chemistry of Macromolecules.” (Wiley, New York, 1961).

83. H. R. Naito, R. Okamoto, T. Sumi, K. Koga, Osmotic second virial coefficients for hydrophobic interactions as a function of solute size. J. Chem. Phys. 156, 221104 (2022).

84. N. A. Perez-Gonzalez, N. D. Rochman, K. Yao, J. Tao, M-T. T. Le, S. Flanary, L. Sablich, B. Toler, E. Crentsil, F. Takaesu, B. Lambrus, J. Huang, V. Fu, P. Chengappa, T. M. Jones, A. J. Holland, S. An, D. Wirtz, R. J. Petrie, K-L. Guan, S. X. Sun, YAP and TAZ Regulate Cell Volume. J. Cell Biol. 218, 3472–3488 (2019).

85. K. Sao, T. M. Jones, A. D. Doyle, D. Maity, G. Schevzov, Y. Chen, P. W. Gunning, R. J. Petrie, Myosin II governs intracellular pressure and traction by distinct tropomyosin-dependent mechanisms. Mol. Biol. Cell 30, 1170–1181 (2019).

86. M. Shahidullah, A. Mandal, R. T. Mathias, J. Gao, D. Križaj, S. Redmon, N. A. Delamere, TRPV1 activation stimulates NKCC1 and increases hydrostatic pressure in the mouse lens. Am. J. Physiol. Cell Physiol. 318, C969–C980 (2020).

87. M. Pallotto, P. V. Watkins, B. Fubara, J. H. Singer, K. L. Briggman, Extracellular space preservation aids the connectomic analysis of neural circuits. eLife 4, e08206 (2015).

88. E. A. Neuwelt, Mechanisms of disease: The blood-brain barrier. Neurosurgery 54, 131–142 (2004).

89. R. Bai, C. S. Springer, D. Plenz, P. J. Basser, Fast, Na+/K+ pump driven, steady-state transcytolemmal water exchange in neuronal tissue: A study of rat brain cortical cultures. Magn. Reson. Med. 79, 3207–3217 (2018). [doi:10.1002/mrm.26980]

90. R. Meli, C. Pirozzi, A. Pelagalli, New perspectives on the potential role of aquaporins (AQPs) in the physiology of inflamation. Front. Physiol. 9, 101 (2018).

91. L. S. King, D. Kozono, P. Agre. From structure to disease: The evolving tale of aquaporin biology. Nat. Rev. Mol. Cell Biol. 5, 687–698 (2004).

92. B. Yang, A. S. Verkman, Water and glycerol permeabilities of aquaporins 1-5 and MIP determined quantitatively by expression of epitope-tagged constructs in Xenopus oocytes. J. Biol. Chem. 272, 16140–16146 (1997).

93. L. Hertz, The glutamate-glutamine (GABA) cycle: Importance of late postnatal development and potential reciprocal interactions between biosynthesis and degration. Front. Endochrinol. 4, 59 (2013).

94. G. W. Koch, E. Schwartz, Isotopic labeling of metabolic water with 18O2. Rapid Comm. Mass Spectrom. 37, e9447 (2023).

95. Q. Kuang, P. Purhonen, H. Hebert, Structure of potassium channels. Cell. Mol. Life Sci. 72, 3677–3693 (2015).

96. R. P. Rand, V. A. Parsegian, D. C. Rau, Intracellular osmotic action. Cell. Mol. Life Sci. 57, 1018–1032 (2000).

97. H. Wiebe, J. Spooner, N. Boon, E. Deglint, E. Edwards, P. Dance, N. and Weinberg, Calculation of molecular volumes and volumes of activation using molecular dynamics simulations. J. Phys. Chem. C 116, 2240–2245 (2012). [10.1021/jp209088u]

98. C. D. Hubbard, R. van Eldik, Mechanistic information on some inorganic and bioinorganic reactions from volume profile analysis. Inorg. Chim. Acta 363, 2357–2374 (2010). [doi: 10.1016/j.ica.2009.09.042]

99. A. Drljaca, C. D. Hubbard, R. van Eldik, T. Asano, M. V. Basilevsky, W. J. le Noble, Activation and reaction volumes in solution. 3. Chem. Revs. 98, 2167–2289 (1998).

